# T cell-intrinsic vitamin A metabolism and its signaling are targets for memory T cell-based cancer immunotherapy

**DOI:** 10.1101/2021.11.19.469290

**Authors:** Fumihiro Fujiki, Soyoko Morimoto, Akiko Katsuhara, Akane Okuda, Saeka Ogawa, Eriko Ueda, Maki Miyazaki, Ayako Isotani, Masahito Ikawa, Sumiyuki Nishida, Hiroko Nakajima, Akihiro Tsuboi, Yoshihiro Oka, Jun Nakata, Naoki Hosen, Atsushi Kumanogoh, Yusuke Oji, Haruo Sugiyama

## Abstract

Memory T cells play an essential role in infectious and tumor immunity. Vitamin A metabolites such as retinoic acid are immune modulators, but the role of vitamin A metabolism in memory T-cell differentiation is unclear. In this study, we identified retinol dehydrogenase 10 (Rdh10), which metabolizes vitamin A to retinal (RAL), as a key molecule for regulating T cell differentiation. T cell-specific Rdh10 deficiency enhanced memory T-cell formation through blocking RAL production in infection model. Epigenetic profiling revealed that retinoic acid receptor (RAR) signaling activated by vitamin A metabolites induced comprehensive epigenetic repression of memory T cell-associated genes, including TCF7, thereby promoting effector T-cell differentiation. Importantly, memory T cells generated by Rdh10 deficiency and blocking RAR signaling elicited potent anti-tumor responses in adoptive T-cell transfer setting. Thus, T cell differentiation is regulated by vitamin A metabolism and its signaling, which should be novel targets for memory T cell-based cancer immunotherapy.

## Introduction

The efficient generation of memory T cells is critical to develop effective cancer immunotherapy and robust protective immunity against infectious diseases. Accumulating evidence demonstrates that nutrient metabolism plays a critical role in T cell differentiation; glycolysis and fatty acid oxidation accelerate effector and memory T cell differentiation, respectively^1–4^. Thus, it may be possible to guide T cells in a favorable direction by manipulating nutrient metabolism^4^.

Vitamin A is an essential nutrient for reproduction, development, cell differentiation, and vision^5–9^. Vitamin A (retinol, ROL) is metabolized into retinoic acid (RA) via retinaldehyde (also known as retinal, RAL) by two oxidation steps that are strictly regulated by retinol dehydrogenases (RDHs) and retinaldehyde dehydrogenases (RALDHs)^9^. Vitamin A and its metabolites are multifunctional and can positively or negatively regulate the acquired immune response. RA, the most active form of these metabolites, enhances the extra-thymic induction of regulatory T (Treg) cells^10, 11^ and is required for the proper function of effector CD8^+^ T cells^12^ as well as the development of both Th1 and Th17 cells^13^. Thus, vitamin A metabolites can influence T cell function and differentiation. However, it remains unsolved whether T cell itself can metabolize vitamin A, and how the metabolites regulate the T cell differentiation.

In this study, we describe that T cell itself can metabolize vitamin A into RAL by RDH10, a rate-limiting enzyme for RA biosynthesis, and that lack of the vitamin A metabolism by T-cell-specific Rdh10 knockout enhances the induction of memory, especially, central memory T cells (T_CM_) in Listeria infection model, indicating that vitamin A metabolites such as RAL and RA regulate T cell differentiation. Furthermore, we describe that RA comprehensively induces repressive chromatin states and deletes the memory T cell profile through retinoic acid receptor (RAR), thereby promoting effector T-cell differentiation. Conversely, blockage of RAR signaling suppresses terminal T cell differentiation and efficiently increases T_CM_ with a potent anti-tumor immunity. Thus, vitamin A metabolism and RAR signaling are novel targets to enhance anti- cancer immunity.

## Results

### Vitamin A metabolism promotes human T cell differentiation

Wilms’ tumor gene 1 (WT1) -specific CD4^+^ T cell clone differentiated during culture into four subsets, CD62L^+^CD127^+^ (P1), CD62L^-^CD127^+^ (P2), CD62L^+^CD127^-^ (P3), and CD62L^-^CD127^-^ (P4) cells, which were considered as central memory (T_CM_), effector memory, effector, and terminal effector T (T_EFF_) cells, respectively, according to cytokine production, proliferation and self-renewal capacity, and pluripotency (Supplementary Fig. 1a–d). Differential gene expression among P1, P2, and P4 was examined using a microarray approach, followed by quantitative PCR. *RDH10* expression increased as a function of the differentiation stage (Supplementary Table 1 and Supplementary Fig. 1e). This was also observed in freshly isolated CD4^+^ and CD8^+^ T cell subsets (Supplementary Fig. 1f). Furthermore, *RDH10* mRNA expression levels in CD4^+^ and CD8^+^ T cells increased sharply and then decreased following T cell receptor (TCR) stimulation (Supplementary Fig. 1g). RDH10 catalyzes the oxidation of ROL to RAL, which is the rate-limiting step in RA biosynthesis^14, 15^. To investigate whether human T cells themselves can metabolize vitamin A via RDH10, RDH10-knocked down and -overexpressed CD4^+^ T cells were generated and examined for RAL and RA production from tritium (^3^H) -labeled ROL. Knockdown (Fig. 1a, b) and overexpression (Fig. 1c, d) of RDH10 decreased and increased the production of the RAL metabolite from ROL, respectively. Importantly, RA, the most active form of vitamin A, was not detected in the activated T cells (Fig. 1b, d).

**Figure 1.**
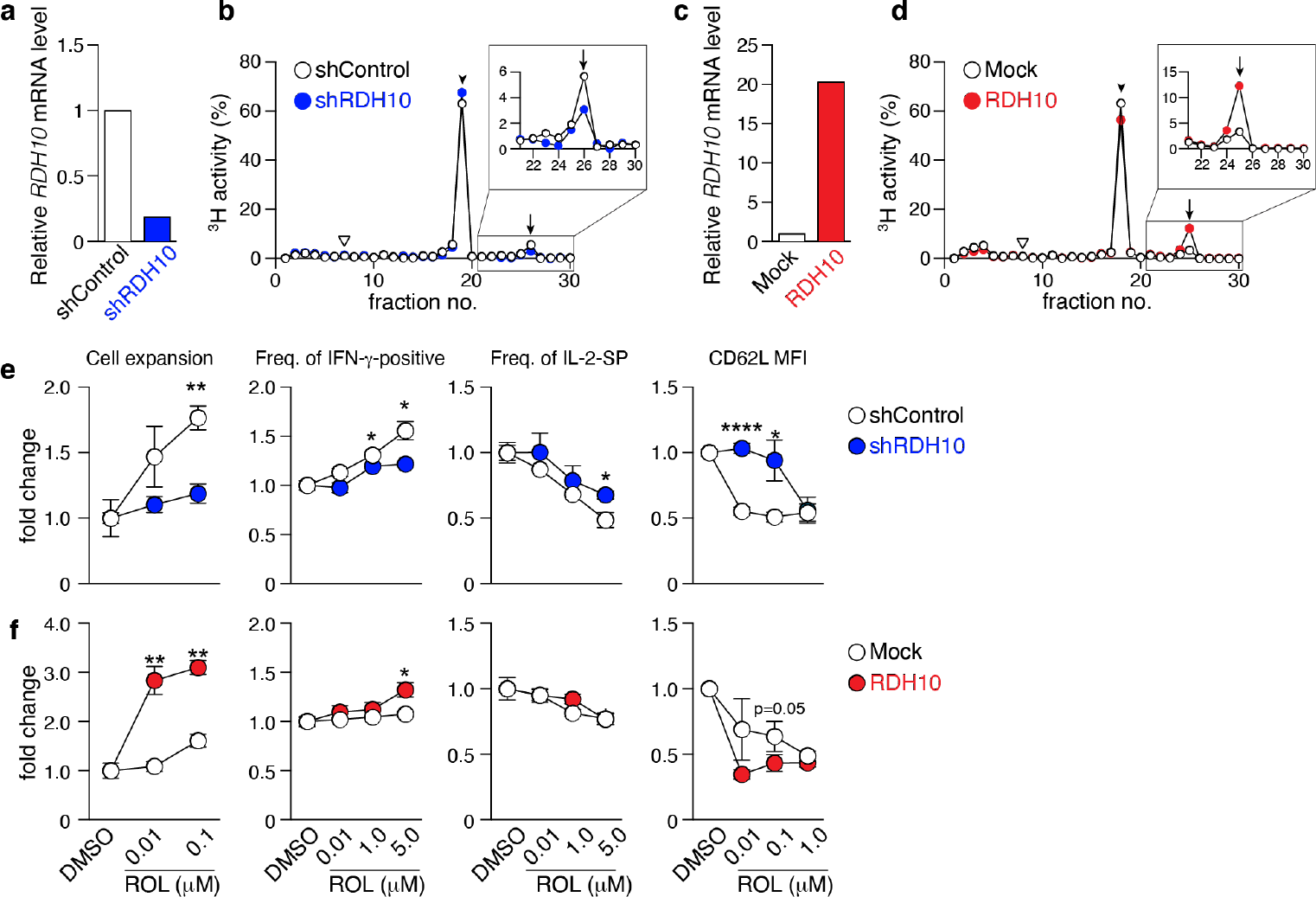
RDH10 metabolizes vitamin A and regulates T cell differentiation. RDH10 in human CD4^+^ T cells was lentivirally knocked down (**a**, **b**, **e**) or overexpressed (**c**, **d**, **f**). **a** and **c** RDH10 mRNA expression level. **b** and **d** shRDH10- (**b**) or RDH10 (**d**)-transduced T cells were cultured in the presence of [^3^H]-all-trans ROL for 4 h. Then, cell extracts were fractionated by HPLC and the radioactivity of each fraction was measured. Arrowhead, arrow, and inverted triangle indicate the peak fraction of standard all-trans ROL, RAL, and RA, respectively. Representative results from two independent experiments are shown. **e** and **f** shRDH10- (**e**) or RDH10 (**f**) -transduced T cells were cultured in the presence of anti-CD3/28 mAbs, IL-2, and all-trans ROL (or DMSO) for 7 days, and then the cell number, cytokine-producing capacity in response to PMA/lonomycin, and expression level of CD62L were measured. Data from three (cell number and cytokine-producing capacity) or five (CD62L expression) independent experiments were analyzed by unpaired t-tests. Error bars show s.e.m. *p < 0.05; **p < 0.01 ; ****p < 0.0001 .

Next, to examine the effect of vitamin A metabolism on T cell differentiation, RDH10- knocked down and -overexpressed CD4^+^ T cells were activated with TCR stimulation and cultured in serum-free medium supplemented with IL-2 and various concentrations of ROL. Seven days later, the cell expansion and phenotypes (CD62L expression and cytokine production capacity) were analyzed. When RDH10 was knocked down, both T cell expansion and IFN-γ-positive T cells decreased, while both IL-2-single positive T cells and CD62L-positive memory phenotype T cells increased, suggesting the blockage of T cell differentiation to terminal stages (Fig. 1e). In contrast, the overexpression of RDH10 resulted in increases in both T cell expansion and IFN-γ- positive T cells, but a decrease in CD62L-positive memory phenotype T cells, suggesting the promoted T cell differentiation (Fig. 1f). The blockage and promotion of T cell differentiation were dependent on the concentration of ROL in serum-free medium without contamination of vitamin A. These results indicate that human T cells themselves metabolize vitamin A, the metabolites of which regulate T cell differentiation, via RDH10.

### Rdh10 and vitamin A regulate CD62L expression in vivo

To examine the roles of Rdh10 in T cell differentiation, T cell-specific Rdh10 conditional knockout (Rdh10CKO) mice were established (Supplementary Fig. 2a). A deficiency of Rdh10 was confirmed by PCR and the loss of RAL production from ROL in T cells (Supplementary Fig. 2b, c). Rdh10-deficient thymocytes exhibited CD4, CD8, CD24, and TCR-β expression levels comparable to those of control mice (Supplementary Fig. 3a). In addition, the frequencies of Foxp3^+^ regulatory and α4β7 integrin-expressing T cells, which are induced by RA ^10, 11, 16^, were not different between Rdh10-deficient and control mice (Supplementary Fig. 3b, c). CD4^+^ and CD8^+^ T cells in lymphoid organs expressed CD62L at higher levels in Rdh10-deficient mice than in control mice (Fig. 2a–c). Enhanced CD62L expression was observed at as early as the double- positive stage when the expression of Cre recombinase was initiated during T cell development in the thymus (Fig. 2d). It is well known that CD62L expression is enhanced in non-activated T cells. However, no difference in the expression of CD44, an activation marker of T cells, between Rdh10-deficient and control mice (Supplementary Fig. 3d) suggested that the enhanced CD62L expression was not due to the change in T cell activation, but was caused by altered vitamin A metabolism. This was consistent with the observation that an Rdh10 deficiency also enhanced CD62L expression in OT-II cells^17^ that are unresponsive to environmental and self-derived antigens in a Rag1-/- background (Fig. 2e).

**Figure 2.**
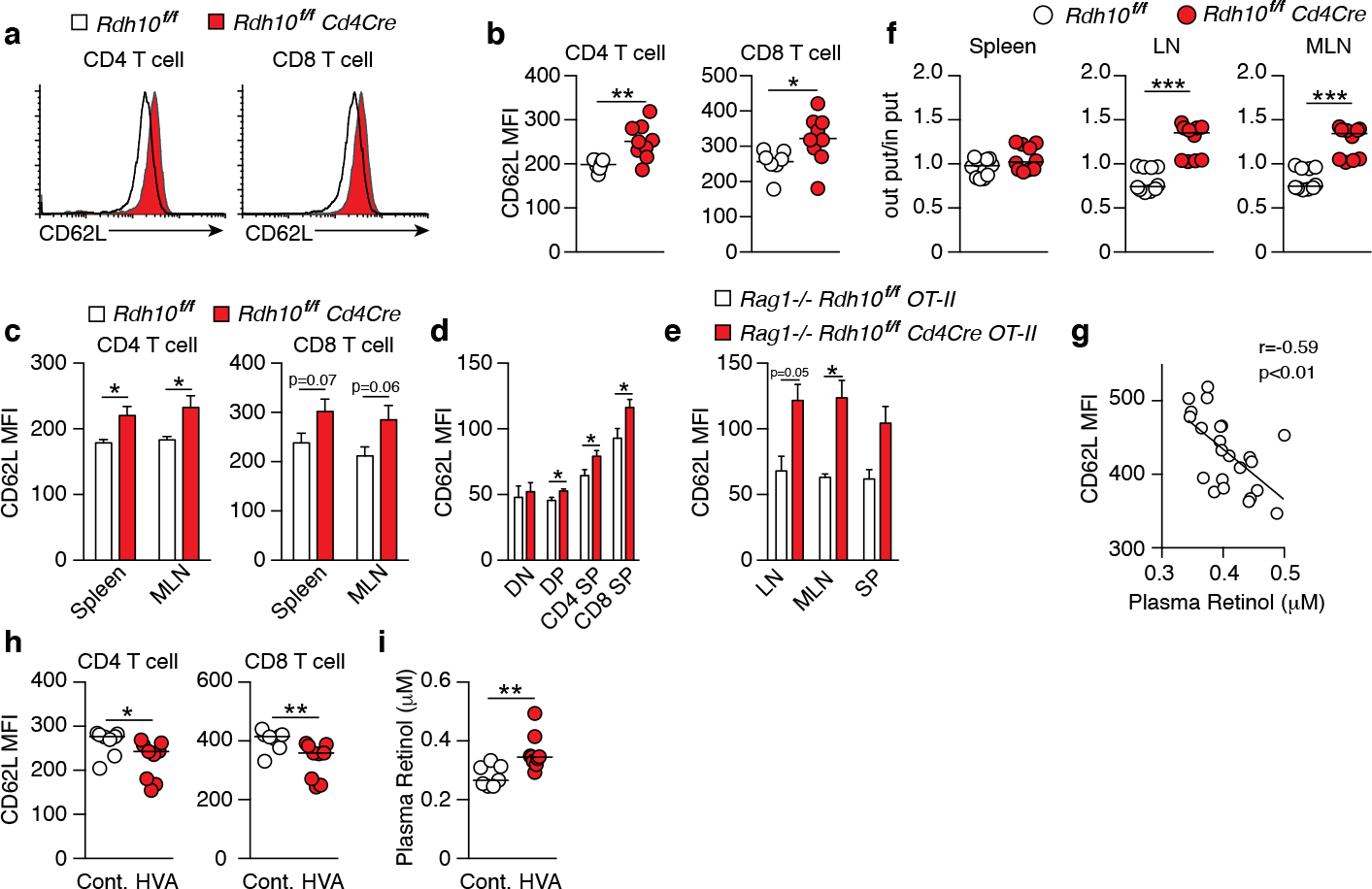
Vitamin A metabolism mediated by Rdh10 physiologically regulates CD62L expression in T cells. **a**–**c** CD62L expression in CD4^+^ and CD8^+^ T cells isolated from the lymph node (LN) (**a** and **b**), spleen, and mesenteric lymph node (MLN) (**c**) of 6- to 8-week-old mice. Data represent n = 7 littermate control and n = 9 Cd4Cre mice. **d** CD62L expression in CD4^-^ CD8^-^ double-negative (DN), CD4^+^ CD8^+^ double-positive (DP), CD4 single-positive (SP), and CD8 SP thymocytes of 5-week-old mice. Data represent n = 8 littermate control and n = 10 Cd4Cre mice. **e** CD62L expression in OT-II cells isolated from the LN, MLN, and spleen (SP) of 6- to 8-week-old mice. Data represent n = 2 littermate control and n = 4 Cd4Cre mice. **f** T cells purified from splenocytes of littermate control and Cd4Cre mice were labeled with either CFSE or CTV, mixed at a ratio of 1:1 and injected into B6 mice (8 × 10^6^ cells per mouse). Donor cells were analyzed in the spleen, LN, and MLN 16 h later. Each symbol indicates one host mouse (n = 10). Data from two independent experiments are shown. **g**–**i** B6 mice were fed a high vitamin A (HVA) or control diet for 1 week. Peripheral blood was collected pre- (**g**) and post-feeding (**h** and **i**), and CD62L expression levels in T cells and the plasma retinol concentration were measured. Data represent CD62L expression levels pre-feeding (n = 23) (**g**) and in control diet-fed (n = 8) and HVA diet-fed (n = 9) mice (**h** and **i**). Data were analyzed by unpaired t-tests (**b**–**f** and **i**), Pearson’s correlation coefficients (**g**), or Mann–Whitney tests (**h**). Error bars show s.e.m. *p < 0.05; **p < 0.01 ; ****p < 0.0001 .

Next, the functional effect of enhanced CD62L expression on T cell homing *in vivo* was examined. The mixtures of Rdh10-deficient and control T cells in the ratio of 1:1 were intravenously injected into B6 mice, and the ratio was investigated in secondary lymphoid organs 16 h later. Consistent with the enhanced expression of CD62L, which was a receptor for T cell migration into lymph nodes. The ratio of Rdh10-deficient T cells to control T cells significantly increased in the lymph nodes (Fig. 2f). These results demonstrated that the ability of Rdh10- deficient T cells to migrate into lymph nodes was potentiated compared to that of control T cells. Thus, these results indicate that Rdh10 regulates the homing capacity of T cells to lymph nodes by the regulation of CD62L expression.

ROL down-regulated CD62L expression in both dose- and Rdh10-dependent manners even in OT-I cells (mouse CD8^+^ T cells) (Supplementary Fig. 4a). Furthermore, RAL and RA, which are ROL metabolites downstream of Rdh10, decreased CD62L expression in OT-I cells independently of Rdh10 (Supplementary Fig. 4b). These results show that ROL metabolism regulates CD62L expression, suggesting that ROL concentration in blood closely links to CD62L expression level on T cells. Indeed, an inverse correlation was observed between plasma ROL concentration and CD62L expression in T cells in wild-type mice (Fig. 2g). These results indicated that food-derived vitamin A can regulate CD62L expression level in T cells. To confirm them, female B6 mice were fed with control diet (4 IU/g) for one week and then with either a high-vitamin A (HVA) (vitamin A, 250 IU/g) or control diet for the next 1 week. As expected, HVA diet reduced CD62L expression in both CD8^+^ and CD4^+^ T cells, with a concomitant increase in ROL concentration in plasma (Fig. 2h, i). Taken together, these results demonstrate that vitamin A (ROL) and its metabolites are physiologically involved in the regulation of CD62L expression in T cells.

### Rdh10 deficiency promotes memory T cell differentiation

The immune response of Rdh10-deficient CD8^+^ T cells was investigated in an ovalbumin-expressing *Listeria monocytogenes* (LM-OVA) infection model ^18, 19^. B6 recipient mice (CD45.2^+^) were transferred with a 1:1 mixture of naïve Rdh10CKO (CD45.1^+^CD45.2^+^) and control (CD45.1^+^CD45.2^-^) OT-I cells, followed by LM-OVA infection. This experimental model made it possible to discriminate the differentiation of Rdh10CKO (CD45.1^+^CD45.2^+^) OT-I cells from that of controls under the same environmental conditions. Upon LM-OVA infection, the proportions of Rdh10CKO OT-I cells were lower at both the effector phase on day 7 and the early contraction phase on day 10 than those of control OT-I cells (Fig. 3a, b). IFN-γ production was lower in Rdh10CKO OT-I cells than in control OT-I cells on day 7 (Supplementary Fig. 5a). Furthermore, short-lived effector cells (SLECs: CD127^lo^KLRG1^hi^) ^20^, which are destined to die at the contraction phase, existed at a low frequency in Rdh10CKO OT-I cells on day 10, whereas memory-precursor effector cells (MPECs: CD127^hi^KLRG1^lo^) ^20^ that would differentiate into long-lived memory T cells were at a high frequency in the OT-I cells (Supplementary Fig. 5b, c). Consequently, the proportion of Rdh10CKO OT-I cells dramatically increased in the memory phase on days 36 and 86, and the proportion of Rdh10CKO OT-I cells became dominant compared to that of control OT-I cells (Fig. 3a–c). Similar kinetics were also observed in the number of transferred OT-I cells (Fig. 3d). These results indicated that an Rdh10 deficiency in OT-I cells induced differentiation toward memory T cells.

**Figure 3.**
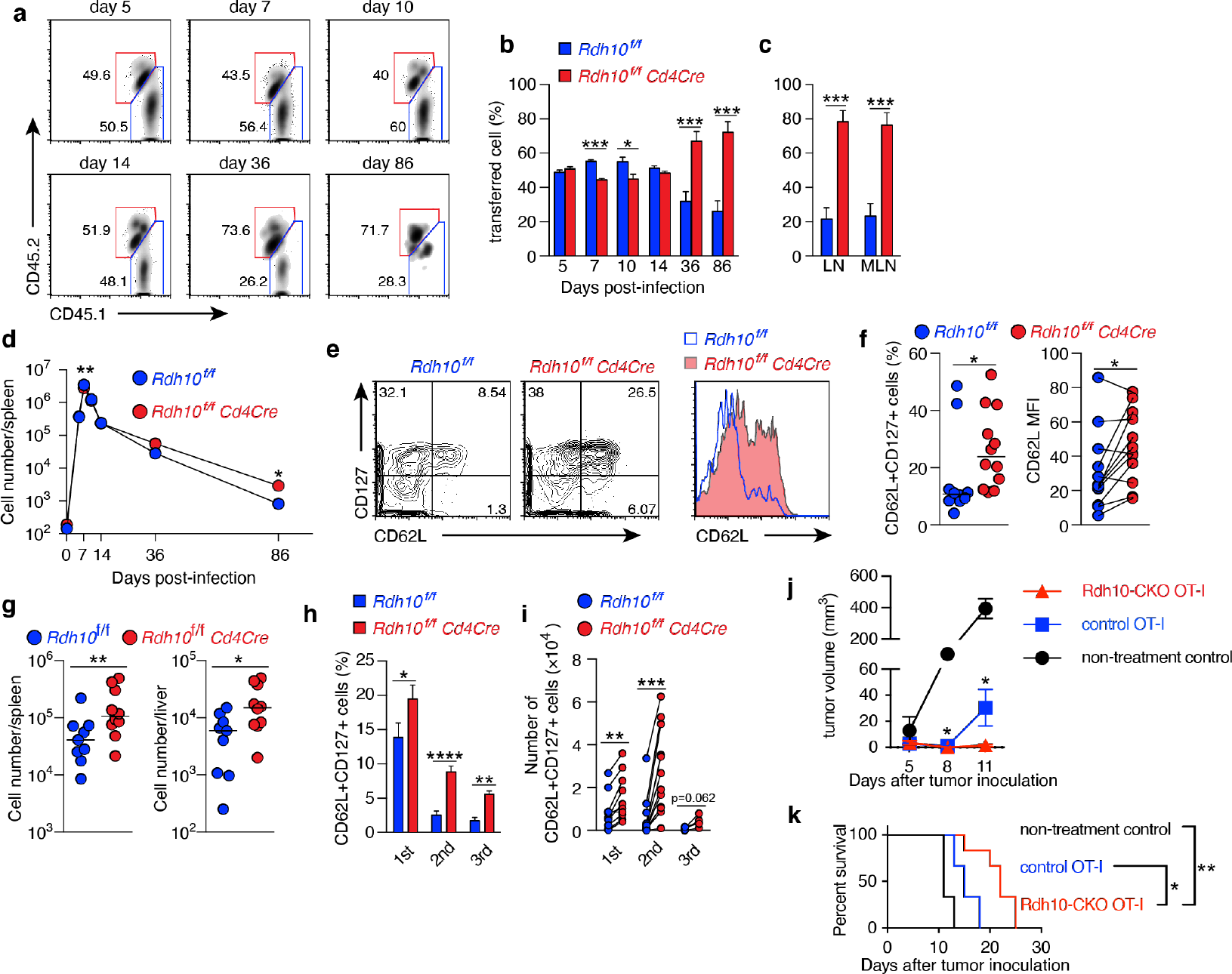
Rdh10 deficiency enhances the generation of central memory T cells. **a-f** OT-I cells were isolated from Rdh1 OCKO (CD45.1’CD45.2’) and control (CD45.1’) mice, mixed at a ratio of 1 :1, and then intravenously transferred into the 86 host (CD45.2’ ). On the next day, the mice were intravenously infected with LM-OVA. **a** Representative dot plots showing the frequency of transferred OT-I cells in the spleen on the indicated days post-infection (pi). band c Frequencies of the indicated OT-I cells among total transferred OT-I cells from the spleen **(b)** and lymph nodes (c, 86 days pi). **d** Kinetics of OT-I cell number in the spleen. Symbols show median values. Data represent n = 5 (day 0), n = 8 (day 5, 7, 10, and 14), and n = 12 (day 36 and 86) mice from two independent experiments **(b-d).** e and **f** Frequency of CD62L’ CD127’ TcM (left) and the level of CD62L expression (right) in OT-I cells from the spleen in the memory phase (>30 days pi). Representative dot plots and histogram (e). Data represent n = 10 (left) or n = 12 (right) control OT-I cells and n = 12 Rdh10CKO OT-I cells **(f). g** Memory OT-I cells were transferred into 86 mice, and OT-I cells were counted in the spleen and liver on day 5 after LM-OVA re-challenge (1 x 106 CFU/mouse). hand **i** Serial transfer of primary (1st) memory OT-I cells and LM-OVA infection were performed. The resultant secondary (2nd) and tertiary (3rd) memory OT-I cells were obtained from the spleen and were measured for the frequency **(h)** and absolute number **(i)** of TcM. Data represent n = 12 (1st and 2nd) and n = 6 (3rd) mice from two independent experiments. **j** and **k** Rdh 1 OCKO or control memory OT-I cells were intravenously transferred immediately after subcutaneous inoculation of EG-7 tumor cells into 3 Gy-irradiated 86 mice. Tumor volumes U) and survival rates **(k)** were evaluated. Data represent n = 3 (non-treatment control and control OT-I) and n = 6 (Rdh10CKO OT-I). Data were analyzed by unpaired !-tests **(b,** c, hand j), Mann-Whitney test **(d,** f-left and **g),** Wilcoxon test (f-right and **i),** or Log-rank test **(k).** Error bars show s.e.m. *p < 0.05; **p < 0.01; ***p < 0.001; ****p < 0.0001.

Next, Rdh10CKO OT-I cells in the memory phase were characterized in more detail. Rdh10CKO OT-I cells expressed CD62L at higher levels and contained T_CM_ (CD62L^+^CD127^+^) cells at a higher frequency in both the spleen (Fig. 3e, f) and lymph nodes (Supplementary Fig. 5d, e), compared to control OT-I cells. The T_CM_ marker CD127, but not the T_EFF_ marker KLRG1, was also expressed at higher levels in Rdh10CKO OT-I cells than in control OT-I cells (Supplementary Fig. 5f, g). To confirm that Rdh10CKO OT-I cells were potential memory T cells, the recall responses of these memory T cells were examined after LM-OVA re-challenge because a strong recall response is a hallmark of T_CM_. A large number of Rdh10CKO OT-I cells were observed in the spleen and liver on day 5 after re-challenge (Fig. 3g). Importantly, these Rdh10CKO OT-I cells were resistant to the attenuation of T_CM_ generation induced by repetitive antigen exposure ^21^ with a higher frequency and a greater absolute T_CM_ count, compared to control OT-I cells (Fig. 3h, i). Thus, these results indicate that Rdh10 deficiency not only enhances memory T cell generation, but also improves the potential for the memory T cells to rapidly expand after re-challenge and to re-generate the T_CM_ pool. The properties of Rdh10-deficient memory T cells suggest Rdh10 as a potential target to improve an effect of cancer immunotherapy because T_CM_ has been shown to have a great anti-tumor activity ^22^. To confirm this concept, Rdh10-deficient memory OT-I cells were investigated for anti-tumor activity against OVA-expressing EL-4 tumor cell line (EG-7) in an adoptive T cell therapy model. As expected, Rdh10-deficient memory OT-I cells suppressed the tumor growth greater than control memory OT-I cells, resulting in prolonged survival (Fig. 3j, k).

In Rdh10 (+/-)-lacZ knock-in reporter mice (Supplementary Fig. 2a), Rdh10 expression was transiently enhanced in OT-I cells after LM-OVA infection (Supplementary Fig. 6a), which was consistent with the transient production of RAL in activated T cells (Supplementary Fig. 6b– d). These findings raised the possibility that RAL produced by Rdh10 in the effector phase could promote the differentiation of T cells into effector T cells. Therefore, the recall response was compared between Rdh10^hi^ and Rdh10^lo^ T cells isolated from Rdh10-lacZ knock-in reporter mice in the effector phase (Supplementary Fig. 7a). Rdh10^lo^ T cells showed a greater recall response than Rdh10^hi^ T cells (Supplementary Fig. 7b, c), indicating that low Rdh10 expression in the effector phase strengthened the recall response as a result of the increase in T_CM_ cells. This was also confirmed by the lower proliferative capacity and lower T_CM_ generation in the recall response in Rdh10-overexpressing T cells (Supplementary Fig. 7d, e). Thus, these results demonstrate that Rdh10 regulates T cell differentiation in the effector phase via RAL.

### RA receptor signaling regulates human T cell differentiation

Not only RA but also RAL binds to retinoic acid receptor (RAR) and the ligand-RAR complex regulates the expression of various genes ^23^. To examine the effect of RAR signaling downstream of RDH10 on human T cell differentiation, CD45RO^+^CD4^+^ T cells were stimulated with anti-CD3 and -CD28 mAbs in the presence of RA or LE540 (an RAR antagonist) ^24^, and the expression levels of T cell differentiation markers (CD62L, CD127, CCR7, and CD27) ^25–27^ were examined after 8 days (Fig. 4a, b, and Supplementary Fig. 8a). RAR signaling promoted by RA decreased T_CM_ phenotype T cells, such as CD62L^+^CD127^+^ and CD62L^+^CD27^+^ T cells, whereas the blockage of RAR signaling by LE540 decreased T_EFF_ phenotype T cells, such as CD62L^-^CD127^-^, CD62L^-^ CCR7^-^, and CD62L^-^CD27^-^ T cells. These results show that the up- and down-regulation of RAR signaling respectively suppress and promote the expression of CD62L indicating that RAR signaling is a key pathway for the regulation of CD62L expression.

**Figure 4.**
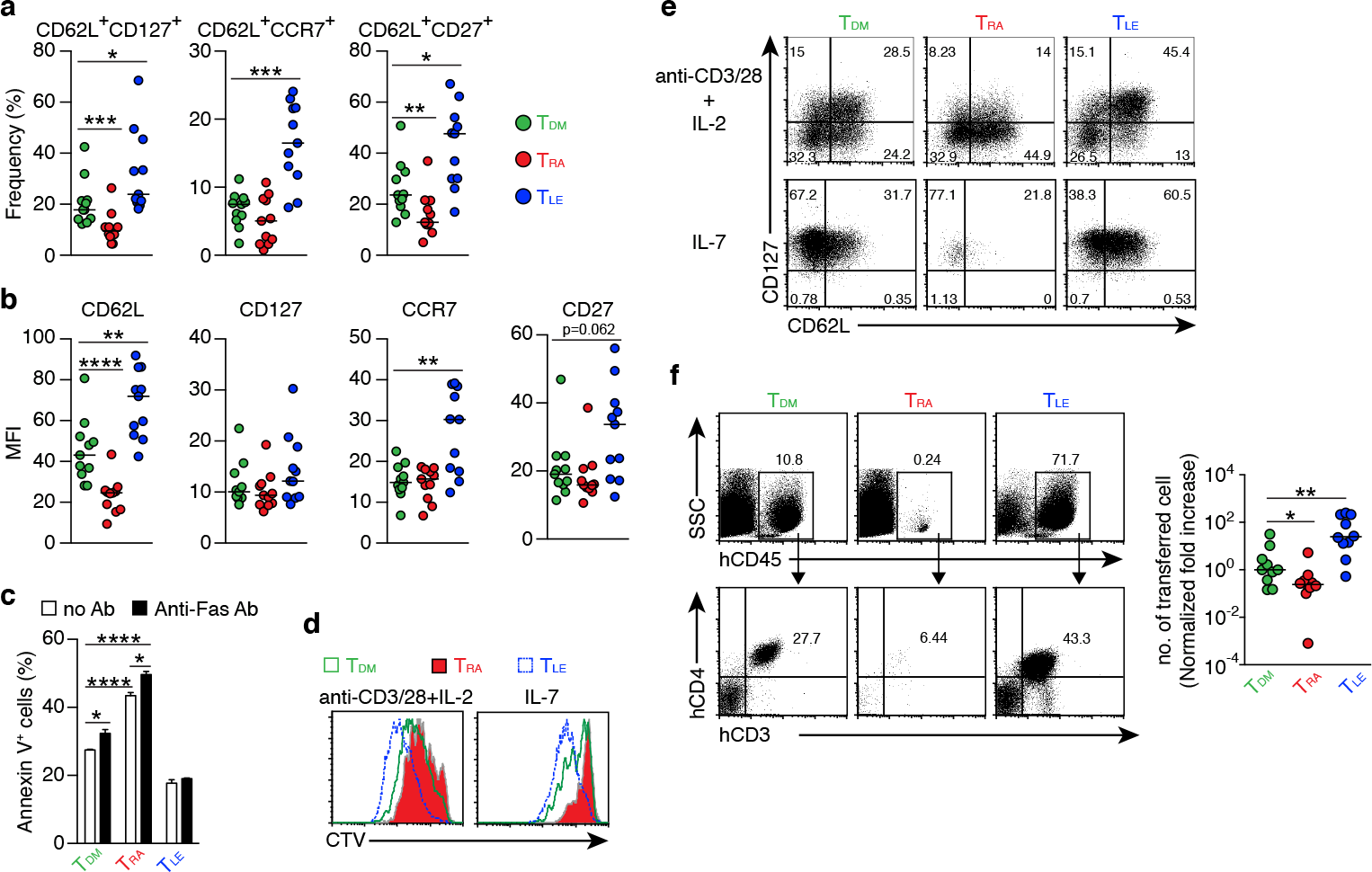
Regulation of T cell differentiation by RAR signaling. Human CD4^+^ CD45RO^+^ T cells were stimulated with anti-CD3/CD28 mAbs in the presence of DMSO, RA (1 μM), or LE540 (10 μM). Seven days later, the T cells were rested in IL-2-free medium overnight and then used for subsequent experiments. **a** and **b** Surface phenotypes of TDM, TRA, and TLE cells. Each symbol indicates the results (n = 11) obtained from six independent experiments. **c** Frequency of apoptotic cells for TDM, TRA, and TLE cells after 4 h of incubation with or without anti-Fas Ab (n = 3). **d** and **e** TDM, TRA, and TLE cells were labeled with CTV and stimulated with the indicated agents. **d** Analysis of cell proliferation using CTV dilution after 4 days of stimulation. **e** The T cells were harvested after 5 days of the stimulation, rested in a cytokine-free medium overnight, and analyzed for their surface phenotypes. Representative data from three independent experiments are shown (**c**-**e**). **f** TDM, TRA, and TLE cells were co-transferred with CD3^+^ T cell-depleted autologous PBMCs into NOG mice. Four weeks later, a flow cytometric analysis was performed to evaluate transferred cells in the spleen. Representative dot plots (Left) and normalized fold increases in the cell number (Right) are shown. The normalized fold increase was calculated by dividing each value by a median value of the corresponding control (i.e., TDM) group. Data from three independent experiments are shown. Each symbol indicates results from a single mouse (n = 10). Data were analyzed by Mann–Whitney tests (**a** and **f**) or unpaired t-tests (**c**). Error bars show s.e.m. *p < 0.05; **p < 0.01; ***p < 0.001; ****p<0.0001.

Next, it was examined whether these RA- and LE540-treated T cells (named as T_RA_ and T_LE_ cells, respectively) exhibited the functional characteristics of T_EFF_ and T_CM_, respectively. DMSO-treated T cells (T_DM_) were used as a control. It is generally known that T_EFF_ is lower in resistance to apoptosis, proliferative capacity, T_CM_ production after cell division, and in vivo expansion capacity, compared to T_CM_. Consistent with these findings, T_RA_ cells showed high frequencies of apoptosis regardless of the presence or absence of anti-Fas mAb (Fig. 4c), and showed decreases in both the proliferative capacity (Fig. 4d and Supplementary Fig. 8b) and the frequency of T_CM_ (CD62L^+^CD127^+^) under two different stimulatory conditions (Fig. 4e and Supplementary Fig. 8c). Furthermore, the number of the transferred T_RA_ cells in NOG mice was lower compared to that of T_DM_ cells 4 weeks after T cell transfer (Fig. 4f). On the other hand, the down-regulation of RAR signaling induced by LE540 had the opposite effect on T cells. Thus, these results demonstrate that RAR signaling promotes T cell differentiation into T_EFF_. It should be noted that the effect of RA on T cell differentiation required TCR signaling (Supplementary Fig. 8d), and that UVI3003^28^, an antagonist of retinoid X receptor (RXR) whose heterodimer with RAR acts as a transcription factor^29, 30^ could not induce T_CM_ (Supplementary Fig. 8e). In addition, RAL down-regulated CD62L expression and decreased T_CM_ generation although these functions of RAL were weaker than those of RA (Supplementary Fig. 8f). Importantly, even in human CD8^+^ T cells, RAR signaling promoted T_EFF_ differentiation as evidenced by low expression of CD62L and weak proliferative capacity in response to IL-7- or TCR-stimulation in RA-treated CD8^+^ T cells (Supplementary Fig. 9).

### Blocking of RAR signaling confers a strong anti-tumor activity on T cells

Anti-tumor activity of T_LE_ cells was examined by using an adoptive T cell therapy model. T cells were transduced with B10-TCR ^31, 32^ which recognized a WT1-derived CTL epitope, WT1_235_ in an HLA-A*24:02-restricted manner, and the T cells were treated with DMSO or LE540. The LE540- treated B10-TCR-transduced T cells (B10-T_LE_ cells) expressed more highly memory or undifferentiated T cell markers such as CD62L, CCR7, and CD45RA, compared to B10-T_DM_ cells (Supplementary Fig. 10). B10-T_DM_ cells or B10-T_LE_ cells were intravenously transferred into K562-A24-luc tumor-bearing NOG mice. In addition, autologous WT1_235_-pulsed, CD3^+^ T cell- depleted PBMCs were co-transferred as antigen-presenting cells into the NOG mouse (Fig. 5a). B10-T_LE_ cells began to suppress tumor growth at day 11 (7 days after T cell transfer) and let the tumor decrease on day 28 (24 days after T cell transfer) whereas statistically significant anti-tumor effect of B10-T_DM_ cells was not observed (Fig. 5b-d). Importantly, B10-TCR^+^ (GFP^+^) CD8^+^ T cells in B10-T_LE_ cells reached a peak on day 18 and were maintained whereas those in B10-T_DM_ cells once reached a peak on day 18 but then remarkably decreased (Fig. 5e). Persistence of tumor-specific CD8^+^ T cells has been shown to be a major factor for eliciting an anti-tumor response in adoptive T-cell therapy ^33, 34^. Therefore, the strong anti-tumor responses of B10-T_LE_ cells likely resulted from the persistence of GFP^+^ CD8^+^ T cells. Thus, these results show that blocking RAR signaling by LE540 confers a potent anti-tumor activity through the increase in memory CD8^+^ T cells that play a central role in anti-tumor immunity.

**Figure 5.**
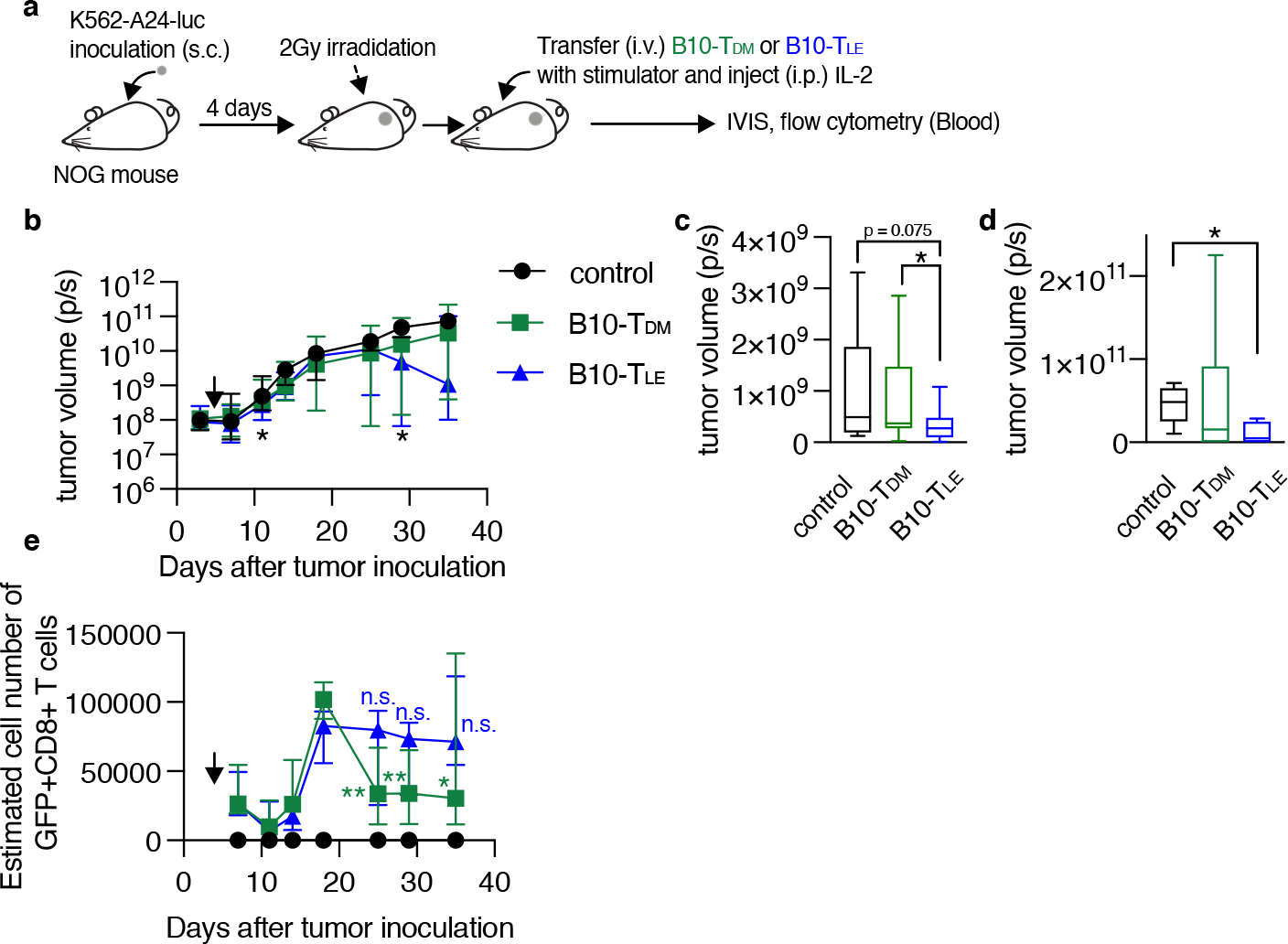
RAR signal blockade confers a strong anti-tumor activity on human T cells. **a** Experimental schematic. B10-TDM or B10-TLE cells were intravenously co-transferred with WT1 peptide-pulsed autologous CD3^+^ cell-depleted PBMCs into K562-A24-luc tumor-bearing NOG mice. Control mice were treated in the same procedure without T-cell transfer. **b** Kinetics of tumor growth. Significant differences were observed in the three groups on days 11 and 29 after tumor inoculation, as marked with asterisk (*). Symbols show median with interquartile range. **c** Tumor volume on day 11 after tumor inoculation (control, n = 11; B10-TDM and B10-TLE, n = 17). **d** Tumor volume on day 29 after tumor inoculation (control and B10-TLE, n = 5; B10-TDM, n = 6). **e** Kinetics of persistence of GFP^+^ CD8^+^ T cells. GFP^+^ CD8^+^ T cells in B10-TDM significantly decreased from a peak (day 18), whereas those in B10-TLE persisted. Symbols show median (with interquartile range) of estimated number of cells (each group, n = 6). The estimated number of cells were calculated from frequencies of GFP^+^ CD8^+^ cells in blood. Data from six (**b** and **c**) or two (**d** and **e**) independent experiments were analyzed by Mann-Whitney test (**c** and **d**) or paired t-test (**e**; day 18 vs. day 25, 29, and 35). *p < 0.05; **p < 0.01.

### Epigenetic regulation of CD62L expression by RAR signaling

CD62L expression in memory T cells is critical for the elimination of tumor cells ^22, 35^ and viral infection ^36^, and the results described above clearly demonstrate that vitamin A metabolism and RAR signaling regulate CD62L expression. Therefore, the mechanism underlying the regulation of CD62L expression was investigated. Enforced expression of RARα decreased CD62L expression at both the protein and mRNA levels in Jurkat cells in the presence of RAL or RA (Fig. 6a, b), indicating that shedding of CD62L from the cell surface ^37^ was not the cause of the down- regulation of surface CD62L expression. KLF2, a key transcription factor that induces CD62L expression ^37–39^, increased regardless of the decrease in CD62L expression by RA treatment (Fig. 6c), suggesting the existence of a novel regulatory mechanism of CD62L expression. RARα directly binds to DNA, recruits co-repressors, and suppresses the transcription of target genes. Both the DNA-binding domain (DBD) and activation function 2 (AF2) domain are critical for regulating target gene transcription ^40, 41^. Neither DBD- nor AF2 domain-deficient RARα proteins down-regulated CD62L expression, regardless of the presence of RA (Fig. 6d). Furthermore, two co-repressor proteins, i.e., nuclear receptor co-repressor (NCoR) ^42^ and silencing mediator for retinoid/thyroid hormone receptor (SMRT) ^43^, were needed for RAR signaling-dependent CD62L repression, as evidenced by significant restoration of CD62L expression after the knockdown of both co-repressors (Fig. 6e). Since these co-repressors recruit histone deacetylases (HDACs) and confer a deacetylated status on histones, the role of histone deacetylation at the *CD62L* locus in the downregulation of CD62L expression was investigated. RA treatment decreased histone acetylation (H3K9/14ac) at the *CD62L* promoter region (Fig. 6f, g and Supplementary Fig. 11). On the other hand, LE540-treatment increased H3K9/14ac (Supplementary Fig. 11). Furthermore, Trichostatin A (TSA), an HDAC inhibitor, could abrogate the RA-induced CD62L downregulation (Fig. 6h). Thus, these results demonstrate that RAR signaling induced by RAL or RA represses CD62L expression via the deacetylation of H3K9/14 at the *CD62L* promoter region.

**Figure 6.**
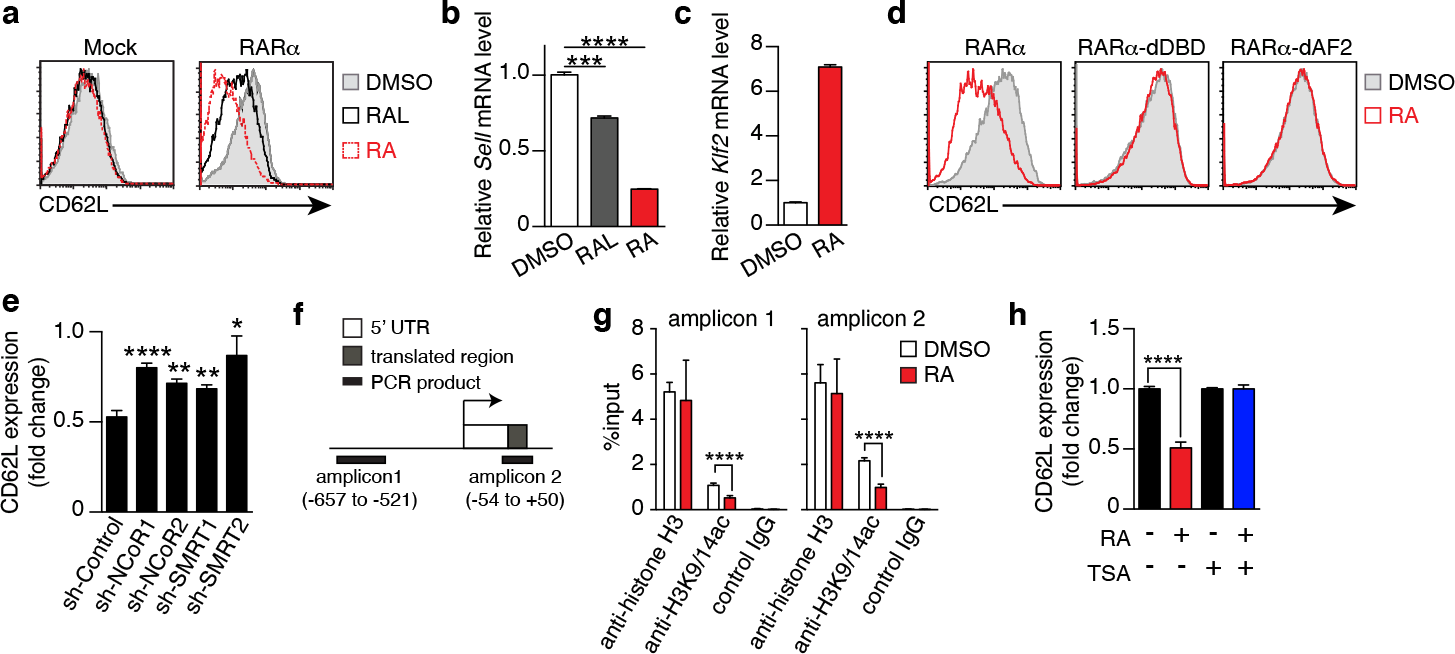
RAR signaling-induced CD62L repression is mediated by epigenetic modifications. **a**–**d** Jurkat cells transduced with the empty vector (Mock), full-length (RARα), DBD-deficient (RARα-dDBD), or AF2-deficient RARα (RARα-dAF2) were cultured for 3–4 days in the presence of RAL (1 μM), RA (1 μM), or DMSO and the expression levels of CD62L protein (**a** and **d**), CD62L mRNA (**b**), and KLF2 mRNA (**c**) were measured. **a** and **d** Representative histograms are shown. **b** and **c** Representative CD62L and KLF2 mRNA levels in RARα-overexpressing Jurkat cells after the indicated treatments. Representative data were from two independent experiments are shown (**a**-**d**). **e** RARα-overexpressing Jurkat cells transduced with the indicated shRNAs were cultured for 4 days in the presence of RA (1 μM) or DMSO, and CD62L protein levels were measured. Data show the fold change of CD62L expression (MFIs of CD62L in RA-treated cells/those in DMSO-treated cells) in three independent experiments. **f** CD62L locus and positions of amplicons for the ChIP assay. **g** ChIP assay results from RARα-overexpressed Jurkat cells treated with DMSO or RA for 4 days. Data were obtained from two (DMSO) or three (RA) independent experiments. **h** CD62L expression (fold change) on RARα-overexpressing Jurkat cells cultured under the indicated conditions (RA, 1 μM; TSA, 50 nM) for 4 days. Data were obtained from three independent experiments. Data were analyzed by unpaired t-tests. Error bars show s.e.m. *p < 0.05; **p < 0.01; ****p < 0.0001.

### RA signaling induces comprehensive epigenetic repression of memory T cell-associated genes

In addition to the down-regulation of CD62L, RA signaling also induced various effector T cell properties, as described in Fig. 4, suggesting that the epigenetic regulation of multiple genes was associated with T cell differentiation. To examine broad-scale epigenetic regulatory effects, a ChIP-seq analysis was performed for T_DM_, T_RA_, and T_LE_ cells with antibodies specific for H3K9/14ac and H3K27me3, which are markers for “opened” and “closed” chromatin structures, respectively^44, 45^. The number of H3K9/14ac-enriched regions was lower in T_RA_ cells (1400 peak calls) than in T_DM_ (2298 peak calls) and T_LE_ cells (2590 peak calls) (Fig. 7a). The number of H3K27me3-enriched regions was comparable among T_DM_, T_RA_, and T_LE_ cells, but was slightly lower in T_LE_ cells than in the other two types of cells. Analysis using the Cis-regulatory Element Annotation System (CEAS)^46^ demonstrated that the proportion of the regions occupied by H3K9/14ac in both the regulatory regions (promoter and downstream) and gene bodies (UTRs, coding exons, and introns) was lower in T_RA_ cells than the other two types of cells (Fig. 7b, upper panel). In contrast, the proportion of the regions occupied by H3K27me3 in these regions, especially in promoter regions was lower in T_LE_ cells than the other two types of cells (Fig. 7b, lower panel). These results revealed that T_RA_ and T_LE_ cells had the “closed” and “opened” chromatin states, respectively, for gene expression.

**Figure 7.**
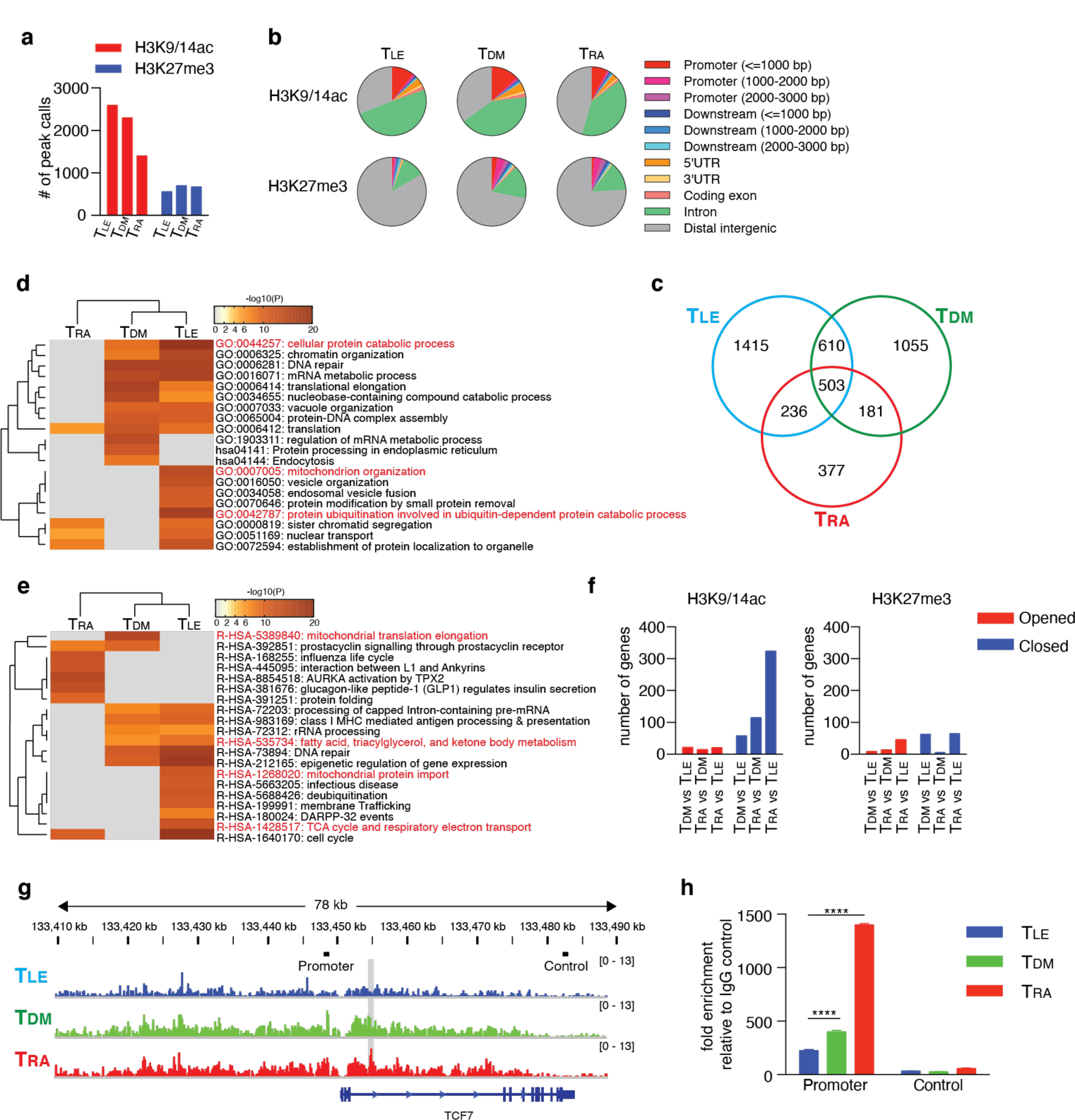
RAR signaling deletes memory T cell profile through comprehensive histone modifications. ChlP-seq using the lllumina HiSeq2500 system was performed with ToM, TRA, and TLE cells for both H3K9/14ac and H3K27me3. a The number of peak calls. b Pie charts show the proportion of H3K9/14ac- and H3K27me3-occupied genomic regions. c Venn diagram shows the number of genes located within ±10 kb of H3K9/14ac-occupied regions. Genes were filtered as follows: p < 1e-5, Fold Enrichment∼ 5, and Peak Tag count∼ 5. d and e Metascape analysis was performed on the uniquely H3K9/14ac-occupied genes (i.e., 1415 genes in TLE, 1055 genes in ToM, and 377 genes in TRA, as described inc). Pathway and Process Enrichment analyses were performed using default settings without Reactome Gene Sets (d) and using only Reactome Gene Sets (e). f Genes located within ±10 kb of H3K9/14ac- and H3K27me3-occupied regions were extracted for the indicated comparison and categorized as closed and opened genes, respectively, during differentiation. The number of closed and opened genes is shown. g The patterns of H3K27me3 peaks at the TCF7 locus are shown. The gray shadow indicates the region that was detected as a peak call in the comparison between TLE and TRA cells. Bars indicated as Promoter or Control are positions of amplicons for ChlP-qPCR (h). h A Ch IP assay was performed for H3K27me3-occupied genes in the naïve CDa• T cells treated with DMSO, RA, or LE540. Representative data are shown from two independent experiments. Data were analyzed by one-way ANOVA with post-hoc tests. Error bars show SD. ****p < 0.0001.

Next, to investigate whether these chromatin states define T cell differentiation, the genes located within 10 kilobases (kb) upstream and downstream of the regions occupied by H3K9/14ac or H3K27me3 were focused. The majority of occupied genes, especially of H3K9/14ac-occupied genes, were different and unique among T_DM_, T_RA_, and T_LE_ cells (Fig. 7c and Supplementary Fig. 12a). A Metascape analysis^47^ of the uniquely occupied genes showed that the cluster of T_LE_ cells was distinct from that of T_RA_ cells, but similar to that of T_DM_ cells (Fig. 7d). T_LE_ cells had, in the uniquely H3K9/14ac-occupied genes, more highly enriched GO terms, including “cellular protein catabolic process”, “protein ubiquitination involved in ubiquitin-dependent protein catabolic process”, and “mitochondrion organization”, which were likely to be associated with memory T cells based on the requirement for autophagy^48^ and mitochondrial oxidative phosphorylation^49^ for memory T cell differentiation, compared to T_DM_ cells (Fig. 7d). Furthermore, an enrichment analysis using Reactome Gene Sets also showed that T_LE_ cells had highly enriched memory-associated genes related to “fatty acid, triacylglycerol, and ketone body metabolism”, “mitochondrial protein import”, and “TCA cycle and respiratory electron transport” (Fig. 7e). Surprisingly, the majority of GO terms for the H3K27me3-occupied genes enriched in T_RA_ and T_DM_ cells were not related to T cell biology, but rather related to the development and differentiation of organs and tissues, such as the heart, eyes, and neurons (Supplementary Fig. 12b). To clarify epigenetic changes among T_DM_, T_RA_ (more differentiated), and T_LE_ cells (less differentiated), the genes with two histone modifications were extracted by a comparison of differentiated T cells with less differentiated T cells (T_DM_ vs T_LE_ cells, T_RA_ vs T_DM_ cells, and T_RA_ vs T_LE_ cells), and categorized into “Opened” and “Closed” genes (Supplementary Tables 2 and 3). This approach also revealed that the most differentiated T_RA_ cells had the pronounced closed chromatin state, compared to T_DM_ and T_LE_ cells (Fig. 7f). Notably, in the comparison of T_LE_ with T_RA_ cells, T_RA_ cells had the closed chromatin state in the regions of memory T cell-associated genes, including *TCF7*, *BCL2*, *KIT*, *DUSP4, CABLES1*, and *SMAD4*^50–53^ (Fig. 7g and Supplementary Table 3). It is well known that TCF7 is a key transcription factor for the maintenance of T cell stemness in CD8^+^ T cells ^54, 55^ and the generation of memory T cells. Consistent with this, H3K27me3 at the *TCF7* locus more increased in RA-treated CD8^+^ T cells compared to LE540- or DMSO-treated CD8^+^ T cells (Fig. 7h). Thus, RA signaling comprehensively induces repressive chromatin states and deletes the memory T cell profile.

## Discussion

Although it was well known that RAR signaling was required for eliciting effector function in both CD4^+^ and CD8^+^ T cells^12, 13, 56, 57^, vitamin A metabolism in T cells and its role for T cell differentiation were unclear. In this study, we revealed that T cells intrinsically metabolized vitamin A to RAL by RDH10 and that this metabolism promoted T cell differentiation into terminal effector T cells. Interestingly, although RDH10 is a rate-limiting enzyme for RA production, T cells could not produce RA, which is a more potent RAR signal activator than RAL. This unique vitamin A metabolism suggests the existence of interesting and important kinetics of T cell differentiation. If T cells can produce RA upon activation, they will rapidly differentiate into terminal T_EFF_ cells and disappear by apoptosis, resulting in the discontinuation of the immune response. On the other hand, RAL not completely but appropriately induces T cell differentiation into T_EFF_ cells to keep T_CM_ cells with the avoidance of the complete T cell terminal differentiation, resulting in the continuation of the immune response. Accordingly, T cells can protect themselves from excessive differentiation by a lack of RA production and maintain a portion of undifferentiated T_CM_ cells. These findings demonstrated a surprising dynamism of T cell differentiation.

Importantly, RA is abundant in tumors ^12, 58^. When T cells infiltrate into tumors, the T cells can take in the exogenous RA that abundantly exists in tumors and rapidly differentiate into T_EFF_ cells. Inflammatory stimuli can also induce RA production from dendritic cells (DCs) ^59^, and thus RA is abundant in inflammatory sites. Therefore, T cells infiltrated into the inflammatory sites can rapidly differentiate into T_EFF_ cells by taking in the exogenous RA and eradicate infectious agents. Moreover, RAL from T cells might be taken up and used for RA synthesis by DCs at inflammatory sites because RA production from DCs is likely to be enhanced by the interaction with T cells^16^. Taken together, while T cells themselves maintain memory function with self-renewal activity and avoid complete differentiation into T_EFF_ cells by a lack of RA production, they can rapidly differentiate into T_EFF_ cells in RA-rich regions such as tumor and inflammatory sites, where prompt effector function is needed.

We demonstrated that RA signaling induced epigenetic repression of memory-associated genes via the decrease and increase in H3K9/14ac and H3K27me3 deposition, respectively. This finding is consistent with a recent report indicating that the accumulation of H3K27me3 deposition on “pro-memory genes”, such as *Tcf7,* guides terminal effector T cell differentiation. In addition, the epigenetic repression by RA signaling also resulted in the disappearance of unique signatures such as the catabolic process and mitochondrial oxidative phosphorylation in memory T cells. At the effector phase, numerous effector T cells against a pathogen can be generated by clonal expansion that is supported by anabolic processes, but most of these cells subsequently die and only ∼5% of them differentiate into memory T cells whose differentiation is supported by catabolic processes, such as autophagy and mitochondrial fatty acid oxidation. Importantly, recent studies have shown that T cells can become memory T cells by shifting their metabolic process from the anabolic to catabolic state^1, 60^, implying that most (95%) effector T cells physiologically fail to change their metabolism to the catabolic state after pathogen clearance. Accordingly, there is surely a “gate pass” for the metabolic change. Our present results indicate that the “gate pass” should be a permissive epigenetic signature, which is turned into a repressive signature by RA signaling.

The mechanisms by which RA signaling induces comprehensive epigenetic repression of memory-associated genes remain unclear. As both the DBD and AF2 domain of RARα are required for the repression of CD62L via the deacetylation of H3K9/14 at the *CD62L* promoter induced by RA, it is likely that RARs bind to genome DNA and recruit functional proteins such as co-repressor proteins, resulting in the induction of comprehensive repression. However, we could not detect any functionally or evolutionarily conserved RA-responsive elements (RAREs) within ±20 kb of the *CD62L* gene (data not shown). In addition, we did not find co-localization of closed regions with RAREs (data not shown). Therefore, it is not clear whether RARs control epigenetic modifications by direct binding to RAREs. It should be noted that RA signaling alone was insufficient for the differentiation into effector T cells, and that TCR signaling was simultaneously required for it, as shown in Supplementary Fig. 8d. Interestingly, upon TCR stimulation, the expression of *Ezh2*, which encodes a methyltransferase, is enhanced and the deposition of H3K27me3 increases in CD8^+^ T cells ^61^. Furthermore, in activated T cells with TCR stimulation, HDACs co-localize with histone acetyltransferases (HATs) and RNA Pol II at transcribed regions to remove acetylation marks and reset the chromatin state^62^. RAR signaling also causes long-term and stable epigenetic changes during the differentiation of stem cells into other specialized cell types, thus silencing or activating large sets of genes either directly or indirectly^63, 64^. Thus, RAR signaling may exhibit cross-talk with TCR signaling in the context of epigenetic modification and achieve the directional differentiation into terminal effector T cells.

Allie et al reported that blockage of RAR signaling enhance the development of central memory-like CD8^+^ T cells in mouse model^57^. However, it was no evidence that blockage of RAR signaling in human T cells enhance anti-tumor immunity through inducing memory T cell. We here succeeded in demonstrating for the first time that RAR signaling in human T cells is targets for improving the clinical efficacy of cancer immunotherapies such as adoptive TCR- or chimeric antigen receptor-transduced T-cell therapy, whose clinical effect is largely dependent upon the amount of memory and undifferentiated T cells^33, 65^. Furthermore, we demonstrated that RDH10- dependent vitamin A metabolism in T cells is also the target. RDH10-targeted immunotherapy should be safe because the conditional RDH10 knockout adult mouse did not present any detectable organ damages (unpublished data). In fact, some chemical agents that could inhibit the RDH10 enzyme activity increased memory T cells (unpublished data). Therefore, the present study contributes to develop memory T cell-based cancer immunotherapy by in vitro and in vivo intervention of T cell differentiation.

## Materials and Methods

### Mice

C57BL/6J mice were purchased from Clea Japan, Inc. (Tokyo, Japan). C57BL/6 mice congenic for the CD45 locus (B6-CD45.1) were purchased from Sankyo Lab Service (Tsukuba, Japan). *Cd4Cre*, *Rag1*^-/-^, OT-I, and OT-II mice were purchased from Taconic (Albany, NY, USA).

NOD/shi-scid/γc^null^ (NOG) mice were obtained from the Central Institute for Experimental Animals (Kawasaki, Kanagawa, Japan). To generate Rdh10 conditional knock-out and Rdh10- lacZ knock-in mice, an *Rdh10* gene targeting vector (PRPGS00072_B_F12) purchased from the International Knockout Mouse Consortium (IKMC) was electroporated into C57BL/6 background embryonic stem cells (EGR-G101). After G418 selection, clones in which homologous recombination correctly occurred were identified by PCR with the following primers: 5′-CAC TAA CTT CTT ACC TTA GTT CAT CCG TC-3′ (GF3) and 5′-CAC AAC GGG TTC TTC TGT TAG TCC-3′ (LAR3) for the 5′ end and 5′-TCT ATA GTC GCA GTA GGC GG-3′ (R2R) and 5′- GCC GGC CGG TCC TGC AAT GGA CTG-3′ (GR3) for the 3′ end. The gene-targeted EGR- G101 was injected into 8-cell stage ICR embryos to obtain chimeric mice. The chimeric males were crossed with C57BL/6J females and germ-line transmission was confirmed by PCR with the following primers: 5′-GCA TTT GTG CTC CCT ACC CAA TCT T-3′ (Rdh5′F) and 5′-CCA ACT GAC CTT GGG CAA GAA CAT-3′ (Common en2R). F1 mice were crossed with CAG-Flpe or CAG-Cre mice to generate Rdh10 floxed (*Rdh10* ^f^) or lacZ knock-in reporter (*Rdh10* ^lacZ^) alleles, respectively (see Extended Data Fig. 3a). Cre recombinase-mediated deletion of the floxed site was confirmed by PCR with the following primers: 5′-TTC ATA AGG CGC ATA ACG ATA CCA C-3′ (P1), 5′-GAA CTG ATC TCA GCC CAG AGA ATA-3′ (P2) and 5′-CCA CCA CCT GAA CAG TGT GGA T-3′ (P3). The *Rdh10*^lacZ^ allele was identified by PCR with 5′-CAC ACC TCC CCC TGA ACC TGA AAC-3′ (RAF5) and P3. All the transgenic and gene-modified mice had C57BL/6 background except for NOG mice. All animals were maintained under specific pathogen-free (SPF) conditions and all animal experiments were approved by the Institutional Animal Care and Use Committee of Osaka University Graduate School of Medicine (approval number 27-009-001). For phenotype analysis of Rdh10-deficient T cells, sex-matched and littermate mice raised in the same cage were used.

### LM-OVA and infection

LM-OVA ^18^ was kindly provided by Dr. H. Shen (University of Pennsylvania, Philadelphia, PA, USA). LM-OVA was prepared as described previously^18, 66^ and injected into the tail vein. Unless otherwise stated, mice were infected with 2 × 10^4^ colony-forming units (CFU) of LM-OVA on the day after adoptive transfer.

### Adoptive transfer and isolation of lymphocytes

For the adoptive transfer of naïve T cells, CD8^+^ T cells (OT-I cells) were isolated from the spleen of naive *Rdh10*^f/f^ *Cd4Cre Rag1*^-/-^ OT-I CD45.1^+^ CD45.2^+^ (Rdh10CKO) or *Rdh10*^f/f^ *Rag1*^-/-^ OT-I CD45.1^+^ (control) mice using the Pan T Cell Isolation Kit II (Miltenyi Biotec, Bergisch Gladbach, Germany). For the adoptive transfer of memory T cells, CD45.1^+^ OT-I cells were isolated from pooled cells of the spleen and lymph nodes from several mice >30 days post-infection by magnetic selection using biotin-conjugated anti-CD45.1 mAb and streptavidin beads. The isolated cells were stained with anti-CD45.2-PE mAb and Rdh10CKO (CD45.1^+^ CD45.2^+^) and control (CD45.1^+^) memory OT-I cells were separately sorted by FACSAria. The isolated Rdh10CKO and control OT-I cells were mixed at a ratio of 1:1 (0.5–1 × 10^4^ each) and then transferred into sex-matched C57BL/6J (CD45.2^+^) mice after the confirmation of the ratio by flow cytometry. For assessment of anti-tumor activity of Rdh10CKO and control memory OT-I cells that were generated in LM-OVA infection model as described above, six-week-old B6 mice were irradiated (3 Gy) and then subcutaneously injected on the left side of abdomen with 2.0 × 10^6^ OVA-expressing EL-4 (EG-7) cells, immediately followed by i.v. injection with 1.5 × 10^5^ memory OT-I cells. In this experiment, we set a humane endpoint; mice were sacrificed when tumor volume reached 15 mm in length. For measurement of tumor volume, blinding was not performed but there were two measurers to obtain accurate data.

### Reagents, antibodies, and flow cytometry

All recombinant interleukins (ILs) were purchased from PeproTec (London, UK), except for human IL-2 (Shionogi, Osaka, Japan). All retinoids (all-trans form), LE540, and UVI3003 were purchased from Sigma-Aldrich (St. Louis, MO, USA), Wako (Osaka, Japan), and Tocris Bioscience (Ellisville, MO, USA), respectively. All antibodies used in this study are listed in Supplementary Table 4. Cells were incubated with FcR blocking Reagent (Miltenyi Biotec) and then stained with the appropriate combinations of mAbs and fluorochrome-conjugated streptavidin. For intracellular cytokine staining, cells were incubated with or without SIINFEKL peptide (1 μM) or PMA + ionomycin in the presence of brefeldin A for 4 h and stained using the Cytofix/Cytoperm Kit (BD Bioscience, Franklin Lakes, NJ, USA). Foxp3-positive cells and apoptotic cells were detected using the BD Foxp3 Staining Kit (BD Bioscience) and Apoptosis Detection Kit (BD Bioscience), respectively. For cell surface-stained samples, 7-aminoactinomycin D (7AAD; eBioscience) was added just before the flow cytometric analysis to exclude dead cells. Flow cytometry was performed using FACSAria. The data were analyzed using FlowJo software (TreeStar, San Carlos, CA, USA).

### Homing assay

T cells were isolated from splenocytes of *Rdh10*^f/f^ *Cd4Cre* and *Rdh10*^f/f^ mice and labeled with either CFSE or CellTrace Violet (CTV). Both labeled cells were mixed at a ratio of 1:1 (4 × 10^6^ per each) and transferred into sex-matched 5–7-week-old B6 mice after the actual ratio was determined by flow cytometry. Sixteen hours later, the ratio of transferred cells from the spleen and lymph nodes was determined by flow cytometry.

### Culture of human cells and assessment of their functions

CD4^+^CD45RO^+^ T cells were freshly isolated from PBMCs of healthy donors using the BD IMag Cell Separation System, stimulated with plate-bound anti-CD3 and soluble anti-CD28 mAbs, and cultured in the presence of IL-2 (20 IU/ml) and RA (1 μM), LE540 (10 μM), or dimethyl sulfoxide (DMSO). Seven days later, the cells were harvested, washed twice, and cultured in an IL-2-free medium overnight. On the next day, the cell phenotypes and functions were evaluated, as described below. To induce apoptosis, the cells were incubated with anti-Fas Ab (100 ng/ml) for 4 h. To assess their proliferation and reconstitution capacity *in vitro*, the cells were labeled with CellTrace Violet (CTV; Thermo Fisher Scientific) and then cultured in the presence of IL-7 (5 ng/ml) or anti- CD3/CD28 mAbs. For reconstitution in NOG mice, the cells (1 × 10^6^) were co-transferred with CD3^+^ cell-depleted autologous PBMCs (2 × 10^6^) isolated using CD3 Microbeads and the LD Column (Miltenyi Biotec) into 5- to 7-week-old female mice. Four weeks later, splenocytes were prepared from the mice and the reconstituted T cells were evaluated by flow cytometry. For assessment of T cell differentiation in RDH10-overexpressed and -silenced T cells, CD4^+^CD45RO^+^ T cells infected with lentivirus vector were purified by sorting and stimulated with anti-CD3/CD28 mAbs in serum-free medium supplemented with the indicated concentration of all-trans ROL and IL-2. Jurkat cells were cultured in the presence of Retinoid (1 μM) or DMSO for 4 days. In treated Jurkat cells, the expression of CD62L was measured at the protein and mRNA levels by flow cytometry and real-time PCR, respectively. All experiments were performed in duplicate or triplicate.

### Anti-tumor activity and persistency of tumor antigen-specific human T cells in NOG mice

B10-T cells were stimulated with plate-bound anti-CD3 (2 μg/ml) and soluble anti-CD28 (2 μg/ml) mAbs, and cultured in the presence of IL-2 (40 IU/ml) and RA (1 μM), LE540 (10 μM), or dimethyl sulfoxide (DMSO). Three days later, the cells were harvested, washed twice, and further cultured for 4 days in 40 IU/ml of IL-2-supplemented medium. For assessment of anti-tumor activity, NOG mice were subcutaneously injected with HLA-A*24:02- and Luciferase gene-expressing K562 leukemia cell line (K562-A24-Luc, 3 × 10^5^ cells). After confirmation of engraftment by In Vivo Imaging System IVIS (Xenogen, Alameda, CA) on day 3, mice were irradiated (2 Gy) and then transferred with LE540- or DMSO-treated B10-T cells (1 × 10^7^ cells) and modified WT1_235_ peptide-pulsed CD3-depleted autologous PBMCs (5 × 10^6^ cells), followed by ip injection of recombinant human IL-2 (500 IU) on day 4. Mice were allocated into groups of equal average base line tumor burden prior to treatments. Tumor volumes were evaluated with IVIS imaging system. In some experiments, blood was collected from tail vein at the same time as that of measurement of tumor volumes and examined for frequency of the transferred GFP^+^ CD8^+^ T cells by using flow cytometry. Cell number of transferred GFP^+^ CD8^+^ T cells in 1 × 10^6^ living cells were estimated with both frequency of transferred T cells (hCD45^+^ hCD3^+^ cells) and frequency of GFP^+^ hCD8^+^ cells in the transferred T cells.

### Preparation and transfection of the viral vector

Human *RDH10* (NM_172037.4) and mouse *Rdh10* (NM_133832.3) cDNA were isolated from PBMCs and splenocytes, respectively. The RARα open reading frame (NP_000955) was sub- cloned from the original plasmid (pFN21AE1591, Promega KK, Tokyo, Japan) into the pFC14K plasmid (Promega). Lentivirus vectors encoding full-length RARα, RARα Δ81-153 (RARα- dDBD), or RARα403 (RARα-dAF2) ^41^ with a C-terminal Halo-tag were generated using the In- Fusion Cloning Kit (TaKaRa Bio Inc, Shiga, Japan). Lentiviruses were generated using CSII-EF-MCS-IRES2-Venus (for overexpression), CS-RfA-EG (for shRNA expression), pCAG-HIVgp, and pCMV-VSVG-RSV-Rev by a standard method. Target sequences for shRNAs are listed in Supplementary Table 5. Retroviruses were generated from the plat-E packaging cell line by the transduction of MSCV-IRES-GFP encoding Rdh10 or from plat-gp packaging cell line by the transduction of MSCV-IRES-GFP encoding B10-TCR and pCMV-VSVG-RSV-Rev. For transfection with the lentivirus, CD4^+^ CD45RO^+^ T cells were stimulated for 3 days as described above and then spin-infected in the presence of polybrene on a RetroNectin-coated plate. Three days later, the cells were maintained in IL-2-free medium overnight, analyzed for their surface phenotype by flow cytometry, and then transfected cells (Venus^+^ cells) were sorted for various experiments, as described above. Lentivirus-transfected Jurkat cells were also generated by a spin-infection method, purified by sorting Venus^+^ or GFP^+^ cells, and then used for experiments. For a generation of B10-T cells, T cells were activated with anti-CD3/CD28 mAbs in the presence of IL-2 for 2-3 days and then spin-infected with B10-TCR-encoding retrovirus as described above. Six hours later, retrovirus infection was repeated. B10-T cells were further cultured for 5-4 days and then treated with RAR agonist/antagonist as described above.

### High-performance liquid chromatography (HPLC)

Naïve and activated T cells (0.5–1.0 × 10^7^ cells) were plated in 3 ml of complete medium supplemented with IL-2 (20 IU/ml) and incubated with 1 μl of ^3^H-labeled ROL (PerkinElmer, Boston, MA, USA) for 4 h. Cells were harvested and washed three times with PBS and intracellular retinoids were extracted as described previously^16^. The extracted retinoids and non-labeled standard all-trans ROL, RAL, and RA were mixed and then fractionated using the HPLC system (Shimadzu, Kyoto, Japan). Separation was performed using the Synergi 4 μm Hydro-RP 80A Column (Phenomenex, Torrance, CA, USA) and an isocratic elution by the solvent composed of 75% acetonitrile and 25% 50 mM ammonium acetate (pH 7) at a flow rate of 1.5 ml/min. Retinoid standards were detected by measuring ultraviolet absorption at 350 nm.

To evaluate the relationship between CD62L expression and plasma ROL, heparinized blood collected from 6- to 7-week-old female mice was used to measure CD62L expression and the concentration of plasma ROL, as described previously^67^. To investigate the effect of a high vitamin A (HVA) diet on CD62L expression, all 6- to 7-week-old female mice were fed an AIN93G-based control (vitamin A, 4 IU/g) diet for 1 week prior to feeding AIN93G-based HVA (250 IU/g). These diets were purchased from Clea Japan, Inc. Extracted retinoids were dissolved in acetonitrile and then analyzed by HPLC as described above. To determine the concentration of ROL, a standard curve was obtained using serial dilutions of all-trans ROL.

### Chromatin immunoprecipitation (ChIP) assay

ChIP assays were performed using a SimpleChIP Plus Enzymatic Chromatin IP Kit (Cell Signaling Technology, Danvers, MA, USA) according to the manufacturer’s instructions.

### Quantitative real-time PCR

mRNAs were extracted using TRIzol reagent (Thermo Scientific) and reverse-transcribed into cDNA using SuperScript VILO (Thermo Scientific). Quantitative real-time PCR was performed using Power SYBR Green Master Mix (Thermo Scientific) following a standard protocol. Primers are listed in Supplementary Table 6. Data were analyzed by a comparative quantification method.

### ChIP-seq analysis

A ChIP-seq and bio-informatics analysis were performed by the TaKaRa Bio Dragon Genomics Center (Yokkaichi, Mie, Japan) using the Illumina HiSeq2500 system. Data quality was checked using FastQC software and all data were found to be of good quality. Reads were aligned to the hg19 human genome using bowtie2 version 2.2.5. Duplicate reads were removed before peak calling using MACS version 1.4.2. Peaks were assigned to genomic regions using BEDTools. According to RefSeq, genes located within 10 kb upstream or downstream of peaks were identified and extracted.

### Statistical analysis

Data were analyzed using Prism 6, 8, or 9. Normally and non-normally distributed data were analyzed by parametric (unpaired *t*-test) and nonparametric (Mann–Whitney test and Wilcoxon test) tests, respectively. For multiple comparisons, ANOVA with post-hoc Dunnett’s test was used. In the case that the objective cell number was less than 10, the samples were excluded from statistical analysis due to unreliable data (Fig. 3f left and Supplementary Fig. 5d).

## Acknowledgments

The authors thank Dr. Hiroyuki Miyoshi (RIKEN Bio-Resource Center, Tsukuba, Japan) for providing the CSII-EF-MCS-IRES2-Venus, CS-RfA-EG, pCAG-HIVgp, and pCMV-VSVG-RSV-Rev plasmid. This study was also supported by Center for Medical Research and Education, Graduate School of Medicine, Osaka University

## Author Contributions

FF and HS designed experiments; FF, SM, AK, AO, SO, EU, MM, AI, MI, SN, HN, AT, YO, JN, NH, and YO performed the experiments; FF, SM, AK, AO, SO, EU, and MM analyzed the data; YO and AK contributed to conception and design of this study. FF and HS wrote the manuscript; all the authors reviewed and approved the final version of the manuscript.

## Data availability

Microarray data and ChIP-seq data will be available through a public database (GSE137142 and GSE137714, respectively). Other data in this study are available from the corresponding author upon reasonable request.

## Appendix

### Supplementary Information

- Supplementary Methods
- Supplementary Figure 1
- Supplementary Figure 2
- Supplementary Figure 3
- Supplementary Figure 4
- Supplementary Figure 5
- Supplementary Figure 6
- Supplementary Figure 7
- Supplementary Figure 8
- Supplementary Figure 9
- Supplementary Figure 10
- Supplementary Figure 11
- Supplementary Figure 12
- Supplementary Table 1
- Supplementary Table 2
- Supplementary Table 3
- Supplementary Table 4
- Supplementary Table 5
- Supplementary Table 6

### Supplementary Methods

#### Housing conditions

Mice were housed in cages (3 or 5 mice per cage), which complied with the Institute of Laboratory Animals Research (ILAR) guidelines, and maintained on MF diet (Oriental Yeast Co., Tokyo, Japan). For the experiments in the infection model, gamma-ray-irradiated CRF-1 diet (Oriental Yeast Co.) were used, instead of MF diet. The animal room had a controlled 12/12-h light/dark cycle (lights on at 8:00 AM), temperature (23 ± 1.5°C), and humidity (45% ± 15%).

#### Microarray

P1, P2, and P4 cells (0.17–1.69 × 10^7^ cells) were sorted from a resting CD4^+^ T-cell clone and RNA was extracted as described above. RNA quality was assessed using the 2100 Bioanalyzer (Agilent Technologies, Santa Clara, CA, USA). A microarray analysis was performed using the Affymetrix GeneChip HG-U133 Plus 2.0 (Affymetrix, Santa Clara, CA, USA) by TaKaRa Bio Inc. Genes were extracted according to the following criteria: 1 ≦ log2 ratio and detection call “Present” in the Experimental sample; -1 ≧ log2 ratio and detection call “Present” in the Base sample. All comparisons fit the two conditions. The extracted genes are listed in Supplementary Table 1. *T-cell assessments using Rdh10-lacZ reporter mouse*

For the detection of Rdh10 expression in T cells and the evaluation of memory formation in Rdh10^hi^ or Rdh10^lo^ T cells, OT-I cells were isolated from the spleen of naïve *Rdh10^lacZ/wt^ Rag1^-/-^* OT-I CD45.1^+^ mice (Rdh10-lacZ reporter mice) and transferred into sex-matched C57BL/6J (CD45.2^+^) mice (1 × 10^5^ per mouse). On the following day, the mice were infected with LM-OVA and then used to detect Rdh10 (lacZ) expression in T cells. For the adoptive transfer of Rdh10^hi^ or Rdh10^lo^ OT-I cells, the transferred OT-I cells were magnetically enriched using biotin-conjugated anti-CD45.1 mAb and streptavidin-conjugated magnetic beads. The enriched cells were stained with fluorescein di-β-d-galactopyranoside (FDG) using the FluoReporter lacZ Flow Cytometry Kit (Thermo Fisher Scientific, San Jose, CA, USA), followed by staining of CD3, CD45.1, and CD45.2. Rdh10^hi^ and Rdh10^lo^ OT-I cells were sorted from the CD3^+^CD45.1^+^CD45.2^-^ fraction and then individually transferred into LM-OVA-infected B6 mice (1 × 10^5^ per mouse). At memory phase, mice were examined for frequencies in the blood of the transferred OT-I cells by flow cytometry and re-challenged with LM-OVA. Five days later, their frequencies were measured again in the blood.

#### In vivo functional assay of Rdh10-overexpressed OT-I cells

CD8^+^ T cells were isolated from splenocytes of naïve OT-I Rag^-/-^ mice, stimulated with irradiated and SIINFEKL peptide-pulsed splenocytes for 20 h, and then spin-infected with a retrovirus encoding mouse Rdh10 or mock control. Twelve hours later, the cells were washed and plated for 48 h in complete medium supplemented with IL-7 (5 ng/ml) and IL-15 (10 ng/ml). GFP^+^ cells were sorted and transferred into wild-type mice (4 × 10^4^ cells/mouse), followed by LM-OVA infection. Subsequently, the expansion and surface phenotype of the transferred GFP^+^ cells were examined in the blood using flow cytometry.

#### Treatment of Human CD8^+^ T cells with RAR agonist/antagonist

Naïve CD8^+^ T cells from peripheral blood mononuclear cells were enriched by Human Naive CD8 T Cell Enrichment Set (BD Bioscience, Franklin Lakes, NJ, USA), stimulated with plate-bound anti-CD3 (2 μg/ml) and soluble anti-CD28 mAbs (2 μg/ml), and cultured in the presence of IL-2 (20 IU/ml) and RA (1 μM), LE540 (10 μM), or dimethyl sulfoxide (DMSO). Three days later, the cells were harvested, washed twice, and further cultured for 4 days in 20 IU/ml of IL-2-supplemented medium. Subsequently, the cells were examined for CD62L expression by flow cytometry or labeled with CTV (1.5 μM) and cultured to investigate proliferative capacity as described elsewhere.

**Supplementary Figure 1.**
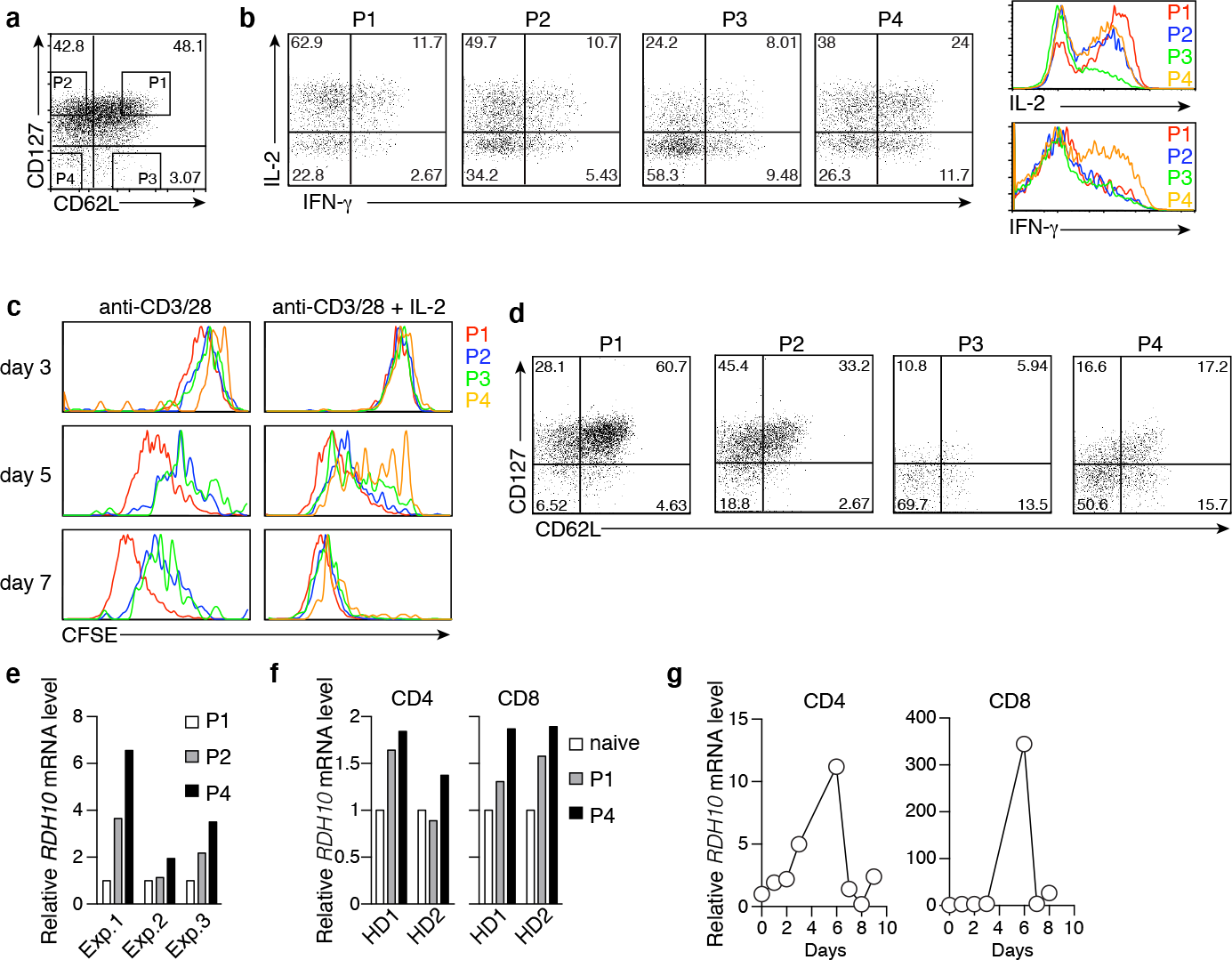
Identification of RDH10 associated with effector T cells. **a** Representative dot plot showing CD62L and CD127 expression in a WT1-specific CD4^+^ T cell clone. **b**-**e** P1–4 populations indicated in **a** were sorted and used for the experiments described below. **b** Representative dot plots and histograms showing cytokine production in each population after 4 h of PMA/Ionomycin stimulation. **c** The T cells were labeled with CFSE, cultured under the indicated conditions, and evaluated for their proliferation capacity. **d** Representative dot plots showing CD62L and CD127 expression after 8 days of culture with antigenic peptide (WT1332)-pulsed dendritic cells. Representative data from three independent experiments are shown (**a**-**d**). **e** RDH10 mRNA expression levels in each population. **f** RDH10 mRNA expression levels in each population from healthy donors (HDs). Each population was sorted from CD4^+^ and CD8^+^ T cells of PBMCs and measured for RDH10 mRNA expression levels. Naïve, CD45RO^-^ CD62L^+^ CD127^+^ T cells; P1, CD45RO^+^ CD62L^+^ CD127^+^ T cells; P4, CD45RO^+^ CD62L^-^ CD127^-^ T cells. **g** Naïve CD4^+^ and CD8^+^ T cells from HD were cultured in the presence of anti-CD3, -CD28 mAb and IL-2. At the indicated time points, RDH10 mRNA expression levels were measured. Representative data from two independent experiments are shown.

**Supplementary Figure 2.**
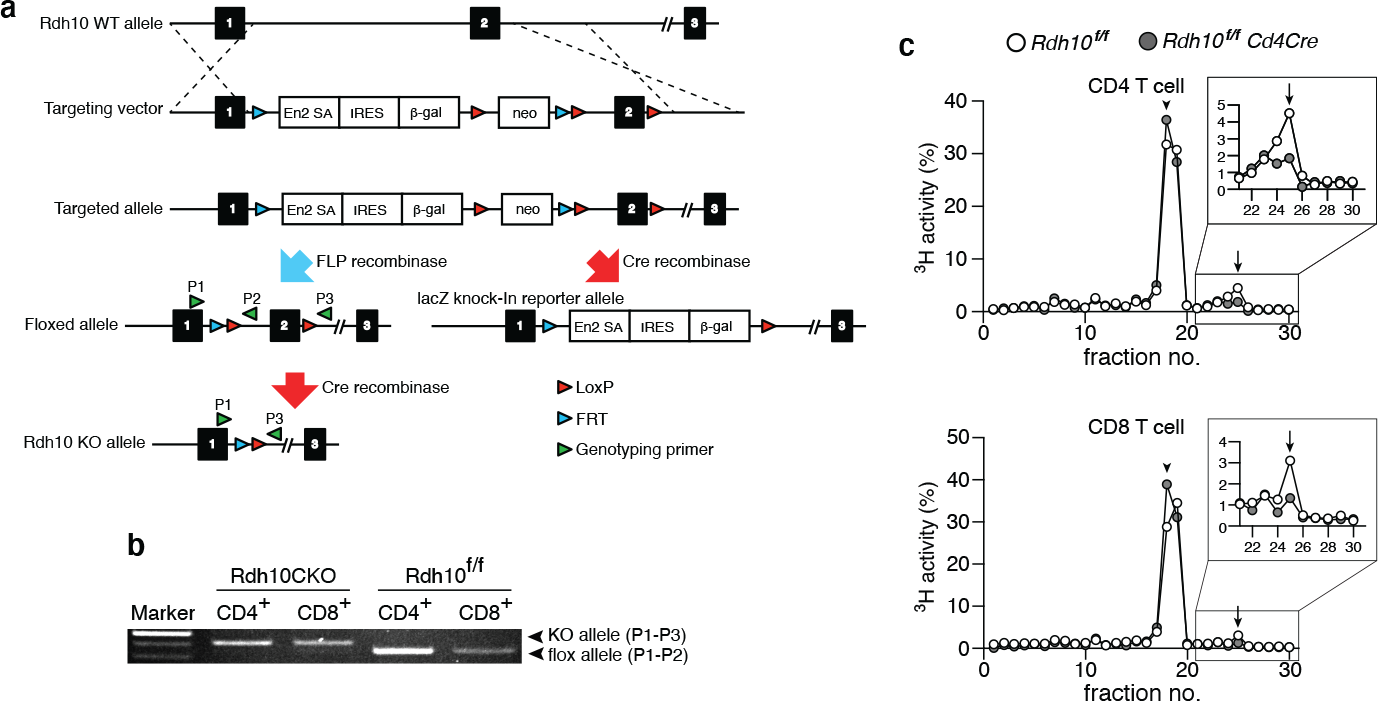
Generation of T cell-specific Rdh10 knockout mice. **a** Schematic of the generation of T cell-specific Rdh10CKO and Rdh10-lacZ reporter mice. **b** Agarose gel shows KO and floxed bands obtained from genotyping using the primers as indicated in **a**. **c** RAL production from ROL was evaluated in Rdh10CKO and control T cells, as described in Fig. 1. CD4^+^ and CD8^+^ T cells were isolated from splenocytes by sorting, expanded for 12 days in the presence of anti-CD3/CD28 mAbs and IL-2, and used for the experiments. The arrowhead and arrow indicate the peak fraction of standard all-trans ROL and RAL, respectively. Representative data from three independent experiments are shown.

**Supplementary Figure 3.**
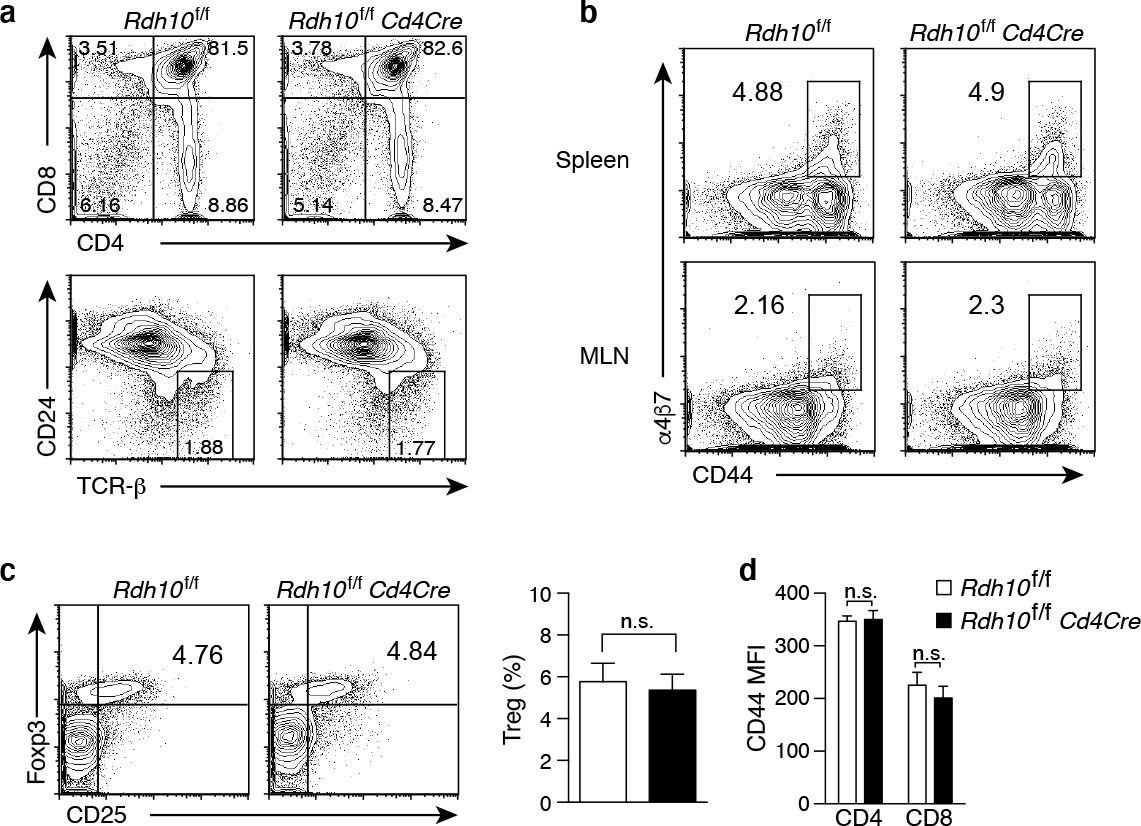
No effect of the loss of Rdh10 on T cell development and Treg and α4β7+ T cell induction. **a** Representative dot plots showing the expression of CD4 and CD8 (*upper*), and TCR-β and CD24 (*lower*) on thymocytes from 4- to 5-week-old mice. **b** Representative dot plots showing the expression of CD44 and α4β7 integrin on CD4^+^ T cells from the spleen and MLN of 6-week-old mice. **c** Frequency of Treg in the MLN from 6- to 8-week-old mice. *left*, Representative dot plots. *right*, Graphs represent n = 8 control and n = 5 Cd4Cre mice. **d** CD44 expression on T cells from splenocytes of 6- to 8-week-old mice. Graphs represent n = 7 littermate control and n = 9 Cd4Cre mice. Data were analyzed by unpaired t-tests (**c** and **d**). Error bars show s.e.m. n.s., not significant.

**Supplementary Figure 4.**
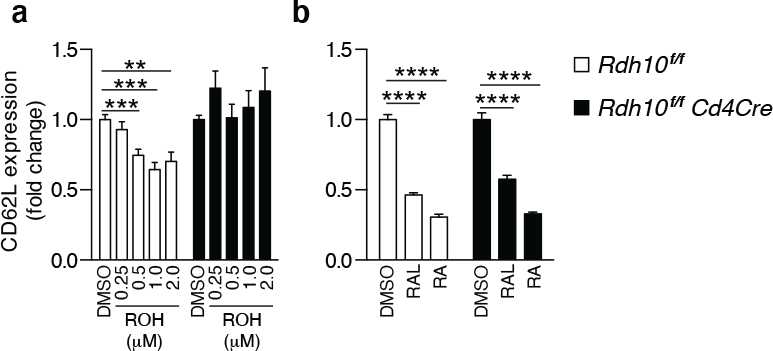
Vitamin A metabolism by Rdh10 regulates CD62L expression in T cells. **a** and **b** OT-I cells isolated from Rdh10CKO and control mice were stimulated with anti-CD3/CD28 mAbs and IL-2 in the presence of ROL, RAL (1 μM), or RA (1 μM). Four days later, CD62L expression in T cells was measured by flow cytometry. Data were obtained from three (**a**) or two (**b**) independent experiments. Data were analyzed by unpaired t-tests. Error bars show s.e.m. **p < 0.01; ***p < 0.001; ****p < 0.0001.

**Supplementary Figure 5.**
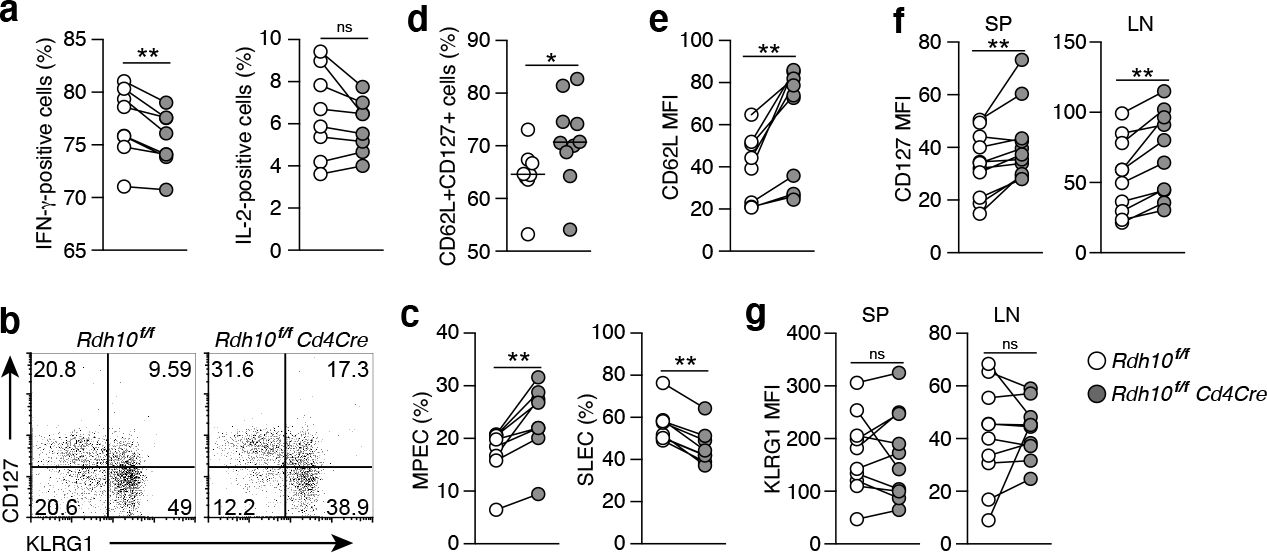
Flow cytometric analysis of Rdh10-deficient OT-I cells in an LM-OVA infection model. **a**-**g** Flow cytometric analysis associated with Fig. 3. **a** Frequencies of IFN-© (left) and IL-2 (right) -producing cells. Splenic OT-I cells from mice on day 7 post-infection (pi) were stimulated with SIINFEKL peptide for 4 h, and analyzed. **b** and **c** Frequencies of MPEC and SLEC in the spleen on day 10 pi. **d** and **e** Frequency of CD62L^+^ CD127^+^ T cells (**d**) and expression levels of CD62L (**e**) in OT-I cells in the memory phase (>30 days pi) in the lymph node. Data represent n = 8 control and n = 10 Rdh10CKO OT-I cells (**d**), and n = 8 pairs (**e**). **f** and **g** Expression of CD127 (**f**) and KLRG1 (**g**) in OT-I cells in the memory phase (>30 days pi) in the spleen and lymph node. Data represent n = 10 pairs. Data were analyzed by Wilcoxon test (**a**, **c** and **e**-**g**) or Mann–Whitney tests (**d**). *p < 0.05; **p < 0.01.

**Supplementary Figure 6.**
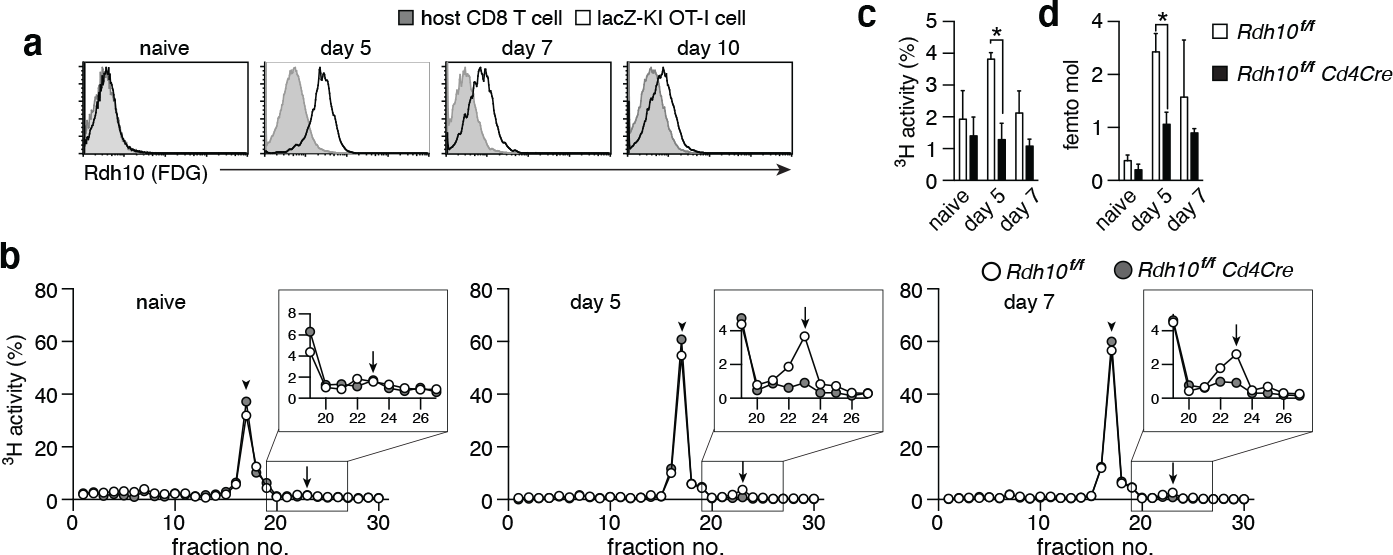
Transient Rdh10 induction and the resultant RAL production in activated T cells. **a** Naïve OT-I cells from heterozygous Rdh10-lacZ knock-in reporter mice were intravenously transferred into B6 mice. On the following day, the mice were infected with LM-OVA via the tail vein. At the indicated time points after infection, lacZ (Rdh10) expression in OT-I cells was evaluated by FDG-staining in the spleen. Representative histograms are shown. **b–d** OT-I cells from Rdh10CKO and control mice were stimulated with anti-CD3/CD28 mAbs and cultured in the presence of IL-2. At the indicated days, the capacity for RAL production in OT-I cells was measured. **b** Representative data from two independent experiments are shown. Arrowhead and arrow indicate the peak fraction of standard all-trans ROL and RAL, respectively. The ^3^H activity (**c**) and absolute amount (**d**) of RAL are shown. Data from two independent experiments were analyzed by unpaired t-tests. Error bars show s.e.m. *p < 0.05.

**Supplementary Figure 7.**
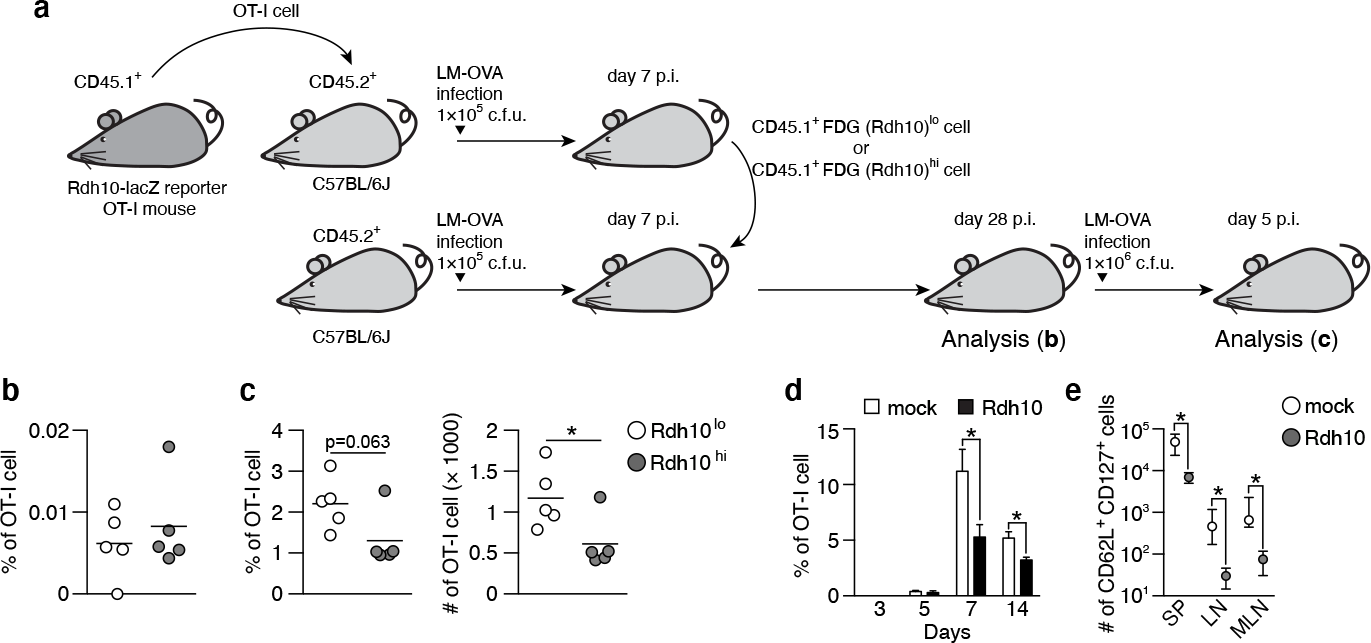
Rdh10 expression at the effector phase dictates the strength of the recall response. **a** Experimental schematic. **b** Frequency of OT-I cells in the blood on day 28 pi. **c** Frequency (*left*) and estimated number (*right*) of OT-I cells in the blood on day 5 after LM-OVA re-challenge. **d** and **e** Freshly isolated OT-I cells were stimulated with SIINFEKL peptide-pulsed splenocytes for 20 h and then retrovirally transduced with Rdh10 or control vector (mock). The transduced OT-I cells were transferred into B6 mice and, on the following day, the mice were infected with LM-OVA. **d** Frequency of OT-I cells at the indicated time points pi. **e** Absolute number of CD62L^+^ CD127^+^ T cells in OT-I cells on day 14 pi. Data represent n=4 Rdh10-overexpressed and n=4 control OT-I cells (**d** and **e**). Data were analyzed by unpaired t-tests (**c** and **d**) or Mann–Whitney tests (**e**). *p < 0.05.

**Supplementary Figure 8.**
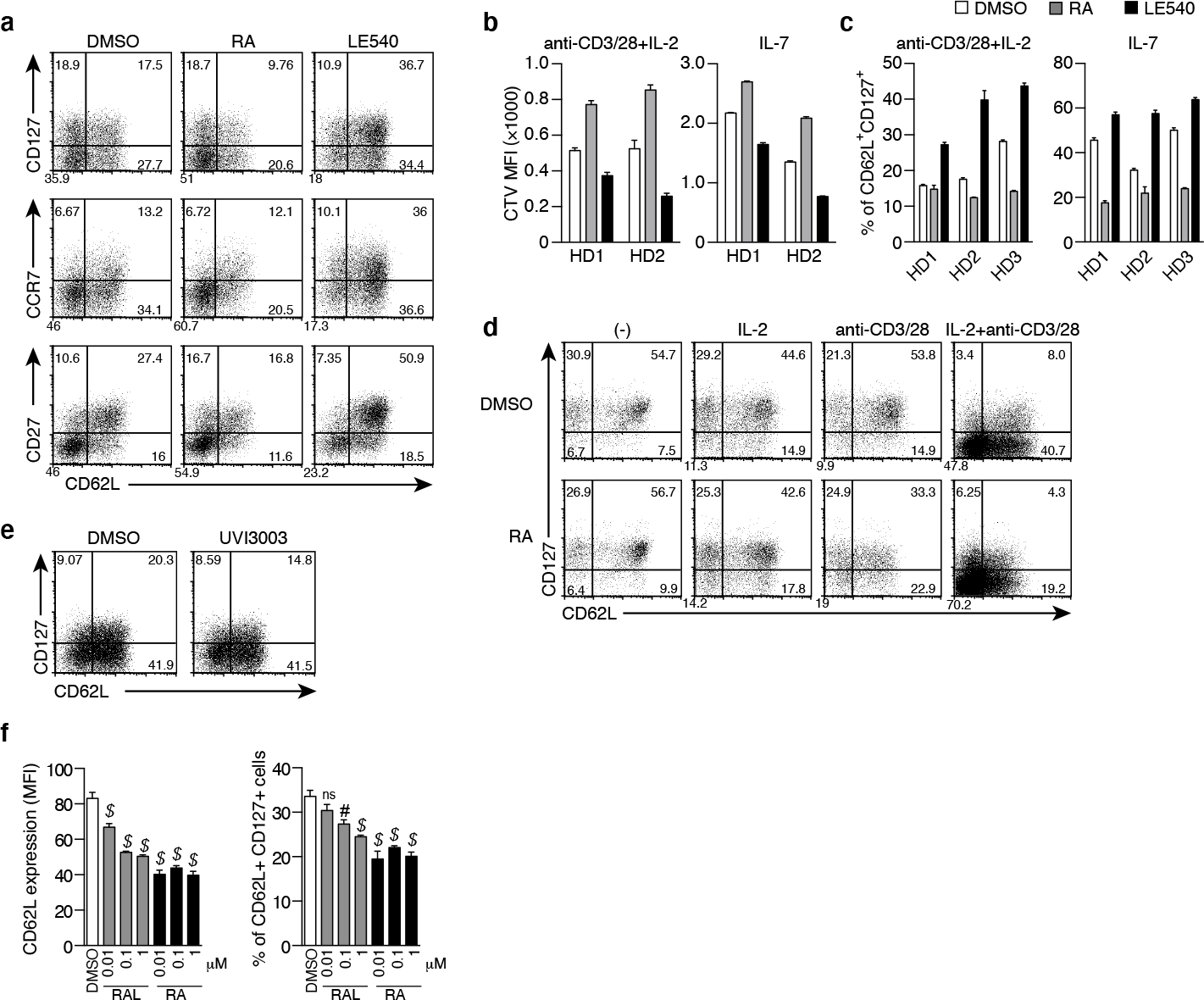
RAR signaling accelerates effector T cell differentiation. CD45RO^+^ CD4^+^ T cells were treated with DMSO, RA, or LE540 as described in Fig. 4 and then the treated T cells were used for the following experiments (**a**–**c**). **a** Representative dot plots showing expression of CD62L, CD127, CCR7, and CD27. **b** and **c** T cells were labeled (**b**) or unlabeled (**c**) with CellTrace Violet (CTV), cultured for 4 or 6 days, respectively, under the indicated conditions, and analyzed for cell proliferation (**b**) and reconstitution capacity of TCM (**c**) by flow cytometry. **d** Representative dot plots showing CD62L and CD127 expression. CD45RO^+^ CD4*^+^* T cells were cultured under the indicated conditions for 7 days and analyzed. **e** Representative dot plots of the T cells treated with a pan-RXR antagonist, UVI3003, as described in Fig. 4. **f** CD45RO^+^ CD4^+^ T cells were treated with either RAL or RA at the indicated concentrations. CD62L expression and frequencies of TCM (CD62L^+^ CD127^+^ cells) were determined by flow cytometry. One-way ANOVA with post-hoc Dunnett’s test was used to compare each treatment to control (DMSO). Error bars, s.em. #p < 0.001; $p < 0.0001.

**Supplementary Figure 9.**
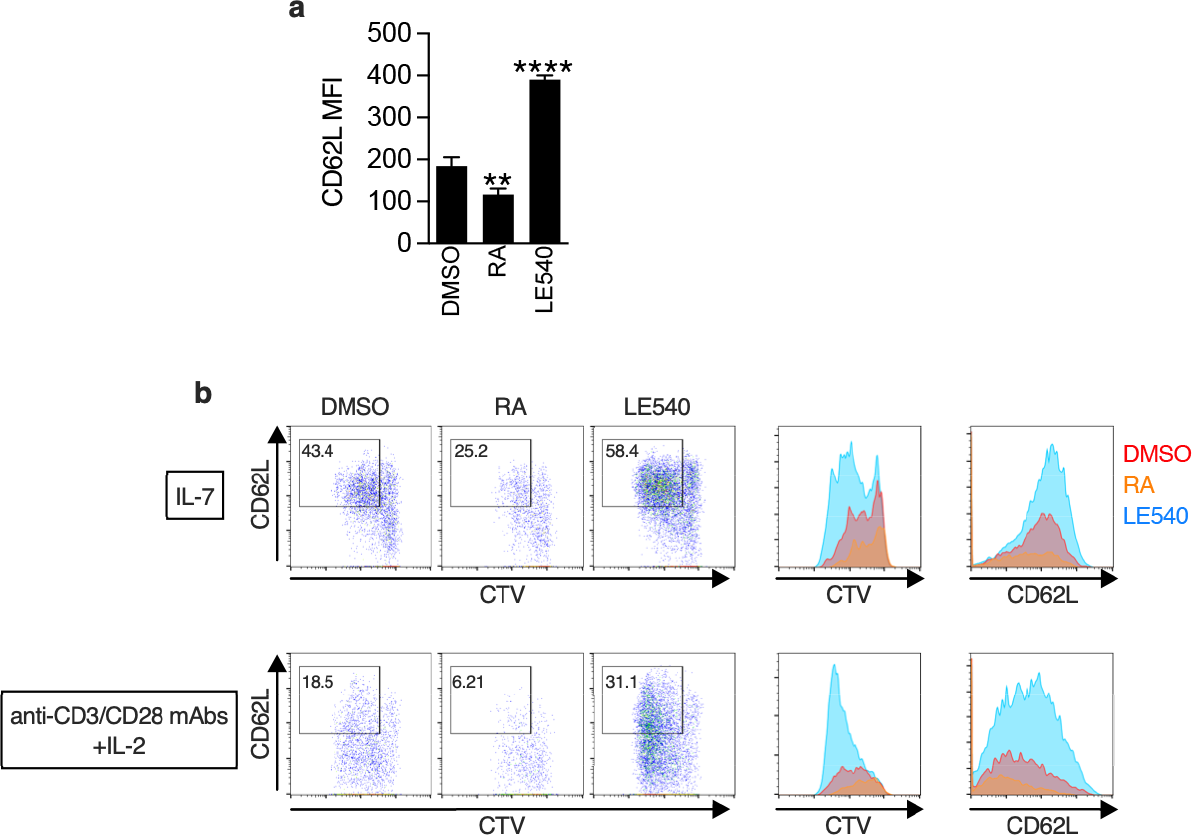
RAR signaling weakens CD62L expression and proliferative capacity in human CD8^+^ T cells. Naive human CD8^+^ T cells were treated with DMSO, RA, or LE540 and then the treated T cells were used for the following experiments. **a** CD62L expression level in the treated T cells. Bars show mean value with SD from triplicate wells. Representative data from two independent experiments are shown. One-way ANOVA with post-hoc Dunnett’s test was used to compare each treatment to control (DMSO). **p < 0.01; ****p < 0.0001. **b** T cells were labeled with CTV, cultured for 3 days under the indicated conditions, and analyzed for cell proliferation and CD62L expression by flow cytometry. The gated cells show CD62L^+^ dividing cells that are indicator of existence of central memory T cells. Representative dot plots and histograms from two independent experiments are shown.

**Supplementary Figure 10.**
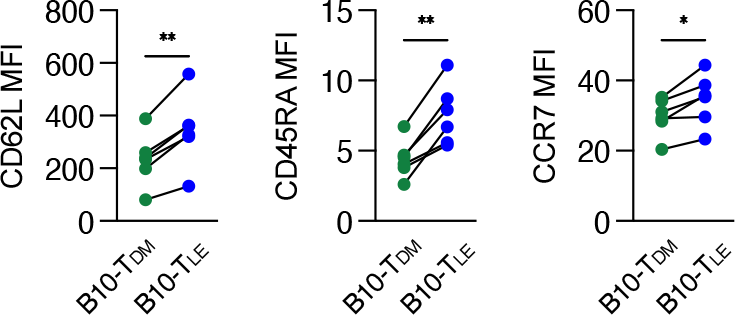
Higher expression of CD62L, CD45RA, and CCR7 in B10-TLE cells. B10-TDM and B10-TLE cells were examined for CD62L, CD45RA, and CCR7 expression level by flow cytometry before adoptive T-cell transfer. The data are from six independent experiments. Paired t-test. *p < 0.05; **p < 0.01.

**Supplementary Figure 11.**
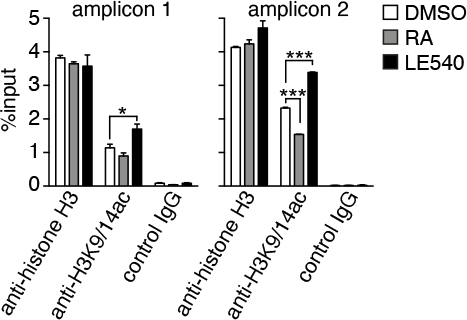
Regulation of the acetylation of histone H3 at the CD62L promoter by RAR signaling. ChIP assays of human CD45RO^+^ T cells treated with DMSO, RA, or LE540 for 7 days as described in Fig 4. Data were obtained from two independent experiments. Data were analyzed by unpaired t-tests. Error bars show s.e.m. *p < 0.05; ***p < 0.001.

**Supplementary Figure 12.**
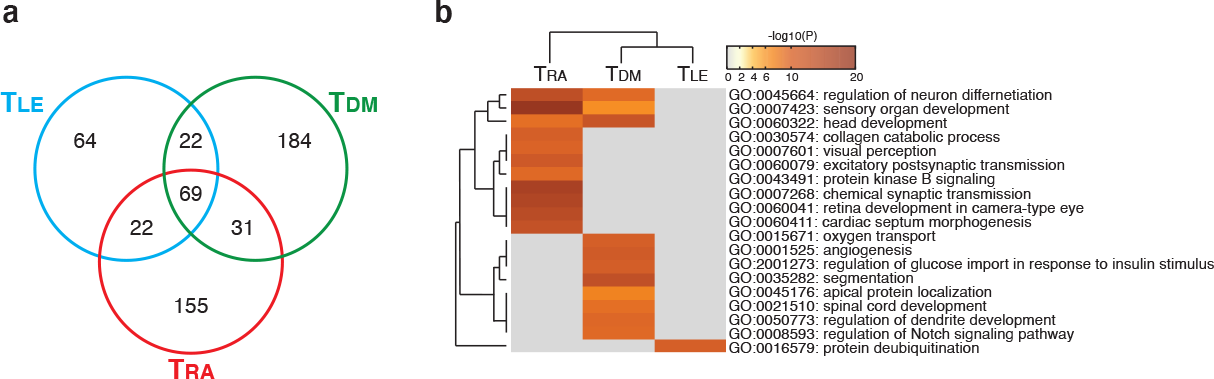
Metascape analysis of uniquely H3K27me3-oocupied genes among T_DM_, T_LE_, and T_RA_ cells. ChIP-seq analysis was performed as described in Fig. 7. **a** Venn diagram shows the number of genes located within ±10 kb of H3K27me3-occupied regions. **b** Metascape analysis was performed on the uniquely H3K27me3-occupied genes. Pathway and Process Enrichment analysis were performed using default settings without Reactome Gene Sets.

